# The Drosophila Y chromosome affects heterochromatin integrity genome-wide

**DOI:** 10.1101/156000

**Authors:** Emily J. Brown, Alison H. Nguyen, Doris Bachtrog

## Abstract

The Drosophila Y-chromosome is gene poor and mainly consists of silenced, repetitive DNA. Nonetheless, the Y influences expression of hundreds of genes genome-wide, possibly by sequestering key components of the heterochromatin machinery away from other positions in the genome. To test the influence of the Y chromosome on the genome-wide chromatin landscape, we assayed the genomic distribution of histone modifications associated with gene activation (H3K4me3), or heterochromatin (H3K9me2 and H3K9me3) in fruit flies with varying sex chromosome complements (X0, XY and XYY males; XX and XXY females). Consistent with the general deficiency of active chromatin modifications on the Y, we find that Y gene dose has little influence on the genomic distribution of H3K4me3. In contrast, both the presence and the number of Y-chromosomes strongly influence genome-wide enrichment patterns of repressive chromatin modifications. Highly repetitive regions such as the pericentromeres, the dot, and the Y chromosome (if present) are enriched for heterochromatic modifications in wildtype males and females, and even more strongly in X0 flies. In contrast, the additional Y chromosome in XYY males and XXY females diminishes the heterochromatic signal in these normally silenced, repeat-rich regions, which is accompanied by an increase in expression of Y-linked repeats. We find hundreds of genes that are expressed differentially between individuals with aberrant sex chromosome karyotypes, many of which also show sex-biased expression in wildtype Drosophila. Thus, Y-chromosomes influence heterochromatin integrity genome-wide, and differences in the chromatin landscape of males and females may also contribute to sex-biased gene expression and sexual dimorphisms.

## Introduction

The Drosophila Y is a degenerated, heterochromatic chromosome with only a few functional genes, primarily specialized in male reproductive function (Carvalho 2002; Carvalho, et al. 2001; Carvalho, et al. 2000; Gatti and Pimpinelli 1983). However, the *D. melanogaster* Y is about 40Mb in size and accounts for ∼20% of the male haploid genome (Gatti and Pimpinelli 1983; Hoskins, et al. 2002) (**Figure 1A**). Most of the Y chromosome is composed of repetitive satellite DNA, transposable elements (TEs), and Y-linked rDNA blocks (Bonaccorsi and Lohe 1991), and it is transcriptionally silenced through heterochromatin formation (Elgin and Reuter 2013). Despite harboring only a few genes, natural variation on the Y chromosome is associated with variation in several traits, including male fitness (Chippindale and Rice 2001) and position effect variegation (PEV), that is, the ability of spreading heterochromatin to induce partial silencing of reporter genes in some cells, resulting in mosaic expression patterns (Gowen and Gay 1934). More recently, it was found that natural variation on the Y has substantial effects on regulation of hundreds of protein-coding genes genome-wide (Dimitri and Pisano 1989; Lemos, et al. 2008; Lemos, et al. 2010; Sackton, et al. 2011).

**Figure 1.**
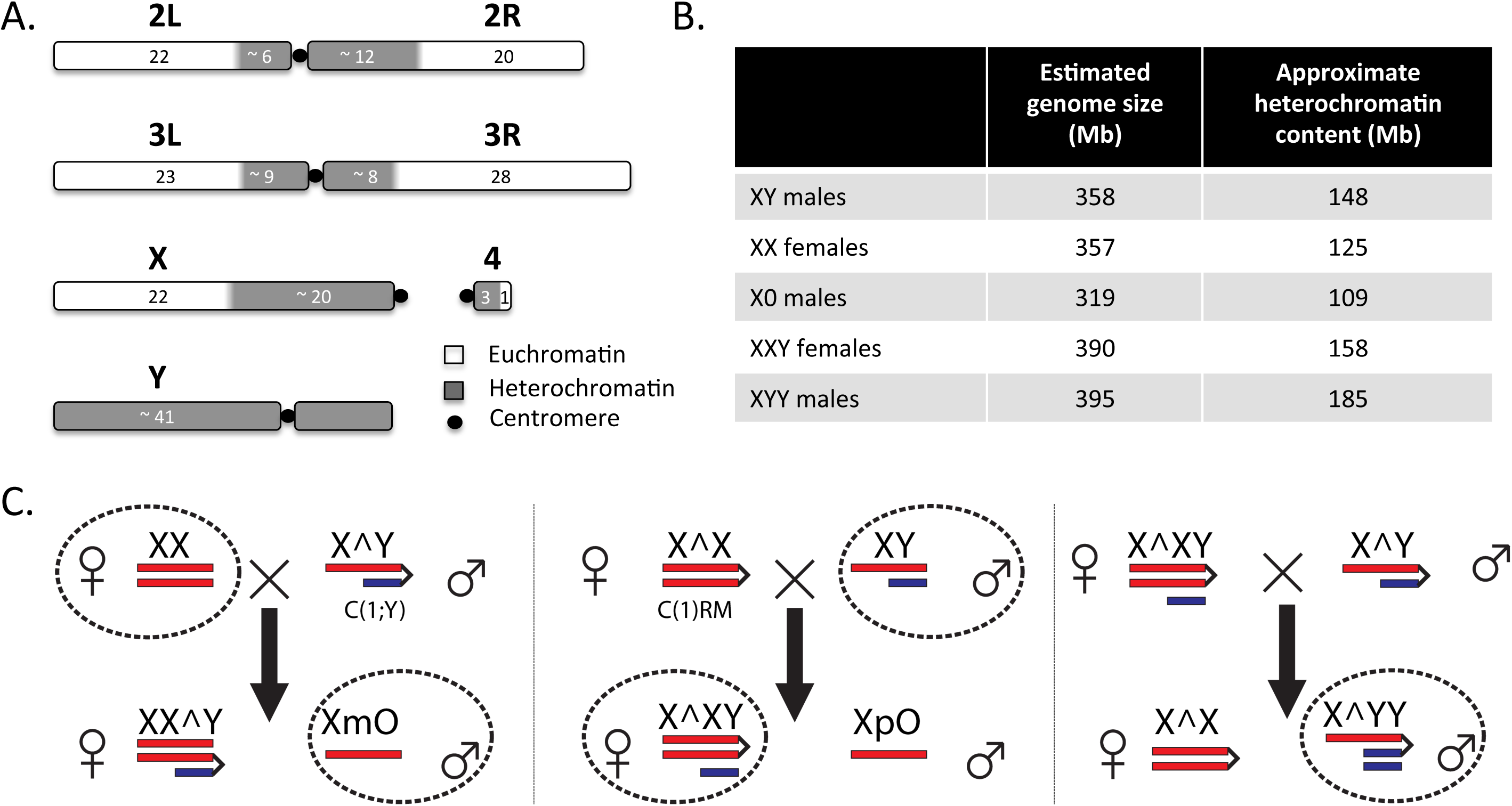
Chromosome structure of *Drosophila melanogaster*, and crossing scheme u3lized. **A.** The le8 and right arms of chromosome 2 (2L, 2R) and 3 (3L, 3R), the small chromosome 4 (the dot chromosome), and the sex chromosomes X and Y are shown (adapted from Hoskins, et al. 2002). The numbers correspond to approximate lengths in megabases, but will differ among Drosophila strains. **B.** Flow cytometry es3mates of the mean diploid genome size of the five karyotypes inves3gated (based on 3 replicate measures). The approximate heterochroma3n content for the strains inves3gated is indicated, assuming that the euchroma3c size is constant for all chromosomes (i.e. 232-Mb for flies with 2 X chromosomes, and 210-Mb for flies with a single X chromosome, see Fig. 1A). **C.** Crossing scheme u3lized to obtain X0 and XYY males, and XXY females (only sex chromosomes are shown). Wildtype Canton-S *D. melanogaster* males and females were crossed to the 2549 strain whose females have C(1)RM and males have C(1;Y). Circled karyotypes were used for the analyses.

The molecular basis of this phenotypic variation is unclear. Single-nucleotide polymorphism in protein-coding genes is low on the Y chromosome (Larracuente and Clark 2013; Zurovcova and Eanes 1999), and it has been proposed that structural variation involving repetitive DNA is responsible for the observed phenotypic effects of different Y chromosomes (Francisco and Lemos 2014). Specifically, most of the highly repetitive Y chromosome is enriched for heterochromatic proteins and repressive histone modifications, and the Y may act as a ‘heterochromatin sink’. That is, the Y chromosome may sequester core components of the heterochromatin machinery (such as structural proteins or modifying enzymes that play key roles in establishing and maintaining heterochromatin), thereby limiting the ability to silence other repetitive regions of the genome (Francisco and Lemos 2014; Henikoff 1996). Under the heterochromatin sink model, Y chromosomes vary in their ability to sequester heterochromatin components due to variations in the total amount or sequence content of their repetitive sequences (i.e. their repeat content). Protein-coding genes from the *D. melanogaster* Y chromosome are only expressed in germ cells of males, but the effects on global gene expression by different Y chromosomes also occur in XXY females and somatic cells of XY males (Lemos, et al. 2008; Lemos, et al. 2010; Sackton, et al. 2011). This observation is consistent with the heterochromatin sink model, where the Y chromosome exerts its effect indirectly by depleting or redistributing chromatin regulators across the genome (Gatti and Pimpinelli 1992). Indeed, PEV assays with different reporter systems have demonstrated that Y-chromosomal DNA suppresses variegation (Dimitri and Pisano 1989; Gowen and Gay 1934). Interestingly, by using a series of cytologically characterized Y chromosome deficiencies and Y fragments, it was shown that Y chromosomes that are cytologically different yet retain similar amounts of heterochromatin are equally effective suppressors, and suppression effect is positively related to the size of the Y-derived DNA (Dimitri and Pisano 1989). This is consistent with the notion that the Y acts as a heterochromatin sink. However, studies to assess the effect of the Y chromosome on heterochromatin formation have been limited to reporter loci through PEV assays (Gowen and Gay 1934), and the global chromatin landscapes of individuals with different amounts of heterochromatic sequence have not yet been directly examined. In particular, studies of PEV do not directly demonstrate changes in the spreading of heterochromatin along the chromosome, but infer it from phenotypic effects on reporter genes (Spofford 1976). In addition, most of the variegating rearrangements have not been characterized at the molecular level and the precise location of their heterochromatic breakpoints has not been determined (Dimitri and Pisano 1989; Gatti and Pimpinelli 1992); it is thus not known whether the different heterochromatic regions are equally effective in inducing variegation. Most importantly, PEV studies do not directly probe the integrity or amount of heterochromatin at repetitive regions that exert PEV through spreading of heterochromatin. Under the heterochromatin sink model, changes in the amount of repetitive DNA should modify the amount of heterochromatin formed at repeats on a global scale. In particular, increasing the amount of repetitive DNA is expected to result in reduced levels of heterochromatin at repetitive regions, since additional repeats should dilute heterochromatic factors that are present in only limited amounts, while decreasing the amount of repetitive DNA should have the opposite effect. Here, we test the hypothesis that the Y chromosome acts to modulate heterochromatin integrity and gene expression genome-wide by contrasting the chromatin landscapes and expression profiles of X0 and XYY males and XXY females to that of wildtype *D. melanogaster*.

## Results

### Fly strains

To compare the chromatin landscape between Drosophila that differ in their sex chromosome karyotype and their amount of repetitive DNA, we set up replicate crosses between *D. melanogaster* stock number 2549 from the Bloomington Stock Center, which has a compound reversed metacentric X chromosome (C(1)RM) or a hetero-compound chromosome with the X chromosome inserted between the two arms of the Y chromosome (C(1;Y)), and the wildtype Canton-S stock (**Figure 1C**). We selected X0 males that contained a maternally transmitted X chromosome (as do wildtype males), and XXY females that contain a wildtype Y chromosome (rather than the C(1;Y) chromosome; see **Figure 1C**). Note that the resulting flies are not isogenic (and it is impossible to create completely isogenic flies using this crossing scheme), but some of the comparisons contrast flies with identical autosomal backgrounds. In particular, our wildtype male and female comparison share the same autosomal genotype (Canton-S), and our X0 males and XXY females both have one autosomal complement from Canton-S, and one from the 2549 stock. XYY males inherit 75% of autosomal genes from strain 2549. We also generated X0, XXY, and XYY flies using a different attached-X and attached X-Y stock, 4248, crossed to the wildtype Canton-S stock; these flies allowed us to verify our findings on chromatin redistribution in an independent genetic background (see below). To get a rough estimate on the amount of repetitive DNA present in the five karyotypes with different sex chromosome configurations, we used flow cytometry to estimate the genome sizes. Under the assumption that the size of the euchromatic chromosome arms is constant across karyotypes, and using estimates of diploid euchromatic genome sizes of 232-Mb for individuals with two X chromosomes and 210-Mb for individuals with one X chromosome (see **Figure 1A**), we estimated the amount of heterochromatic sequences in each karyotype. As expected, we found a gradient of heterochromatic sequence content per diploid cell for the five karyotypes, with XO males (∼109 Mb) < XX females (∼125 Mb) < XY males (∼148 Mb) < XXY females (∼158 Mb) < XYY males (∼185 Mb) (**Table S1**, **Figure 1B**).

### Quantification of histone modifications

We aged independent replicates of all flies for 8 days, and carried out chromatin immunoprecipitation followed by DNA sequencing (ChIP-seq) on head and thorax tissue using commercial antibodies against three post-translational histone modifications (H3K4me3, H3K9me2, H3K9me3). GC content bias, that is, the dependency of read coverage and GC content found in Illumina sequencing data, can be especially problematic for repetitive DNA analysis since repeated sequences often have extreme GC contents. We corrected for GC content biases in our ChIP-seq experiments using a method developed by (Benjamini and Speed 2012) and implemented by (Flynn, et al. 2017). We employed a previously described normalization strategy (Li, et al. 2014) to compare the genomic distribution and relative levels of chromatin marks across flies with different karyotypes. Specifically, we ‘spiked in’ a fixed amount of chromatin from female 3^rd^ instar *Drosophila miranda* to each *D. melanogaster* chromatin sample prior to ChIP and sequencing. *D. miranda* chromatin served as an internal standard for the immunoprecipitation experiment (**Table S2**), and the relative recovery of *D. melanogaster* ChIP signal vs. *D. miranda* ChIP signal, normalized by their respective input counts, was used to quantity the relative abundance of the chromatin mark in *D. melanogaster* (see Methods for details; Li, et al. 2014). Note that this normalization strategy uses input coverage to account for differences in ploidy levels of sex chromosomes among the different karyotypes investigated and is agnostic to the total genome size of the sample (**Figure S1**). *D. miranda* is sufficiently diverged from *D. melanogaster* for sequencing reads to be unambiguously assigned to the correct species: even in repetitive regions, <4% of the reads cross-mapped between species; these regions were excluded from the analysis.

We also used a different normalization strategy to quantify the absolute abundance of the heterochromatic chromatin marks for each *D. melanogaster* karyotype. In particular, the relative recovery of *D. melanogaster* ChIP signal vs. *D. miranda* ChIP signal, normalized by their respective input counts, was estimated using a linear regression model (Bonhoure, et al. 2014, see Methods). Overall enrichment patterns and differences among karyotypes are quantitatively similar between the two methods, showing that our inferences are robust to our normalization strategy (**Figure S2A**). Repetitive regions pose a challenge for mapping with short reads, since one cannot be sure that a particular locus is generating the reads in question if they map to multiple positions. Our study is concerned with the overall behavior of repetitive regions in the genome, and not focused on any particular locus; thus, analyzing all reads (including those mapping to multiple locations) is most appropriate for our purpose. However, we repeated our analysis using only uniquely mapping reads, to confirm that our results are robust to uncertainly in alignments (**Figure S2B**).

Signal for H3K4me3 is highly correlated across samples (**Table S3**), showing that our ChIP data are of high quality. In addition, H3K9me2 and H3K9me3 are known to have very similar genomic distributions (Roy, et al. 2010), and they correlate well with each other for all samples (**Table S3**), and also with independent biological replicate ChIP data without a *D. miranda* chromatin spike (**Table S3**, see Methods for details). Finally, we also generated replicate ChIP-seq data for H3K9me3 from XO, XXY, and XYY individuals using a different attached X stock, 4248. Again, these data are highly correlated and show similar genomic distributions and overall differences among the sex chromosome karyotypes as obtained for the 2549 strain (see below). Thus, our ChIP data are of high quality, and our results are reproducible using different mapping and normalization strategies, and across different histone modifications, independent biological replicates, and different genotypes. We used the total normalized number of *D. melanogaster* reads to compare the genome-wide distribution of chromatin modifications in flies with different sex chromosome karyotypes. **Figure 2** shows the genomic distribution of the active H3K4me3 chromatin mark for the various karyotypes, and **Figures 3** and 4 show genomic distributions for the repressive H3K9me2/3 marks, respectively, at heterochromatic regions, and across the heterochromatin / euchromatin boundary (i.e. the transition of pericentromeric heterochromatin to euchromatin).

**Figure 2.**
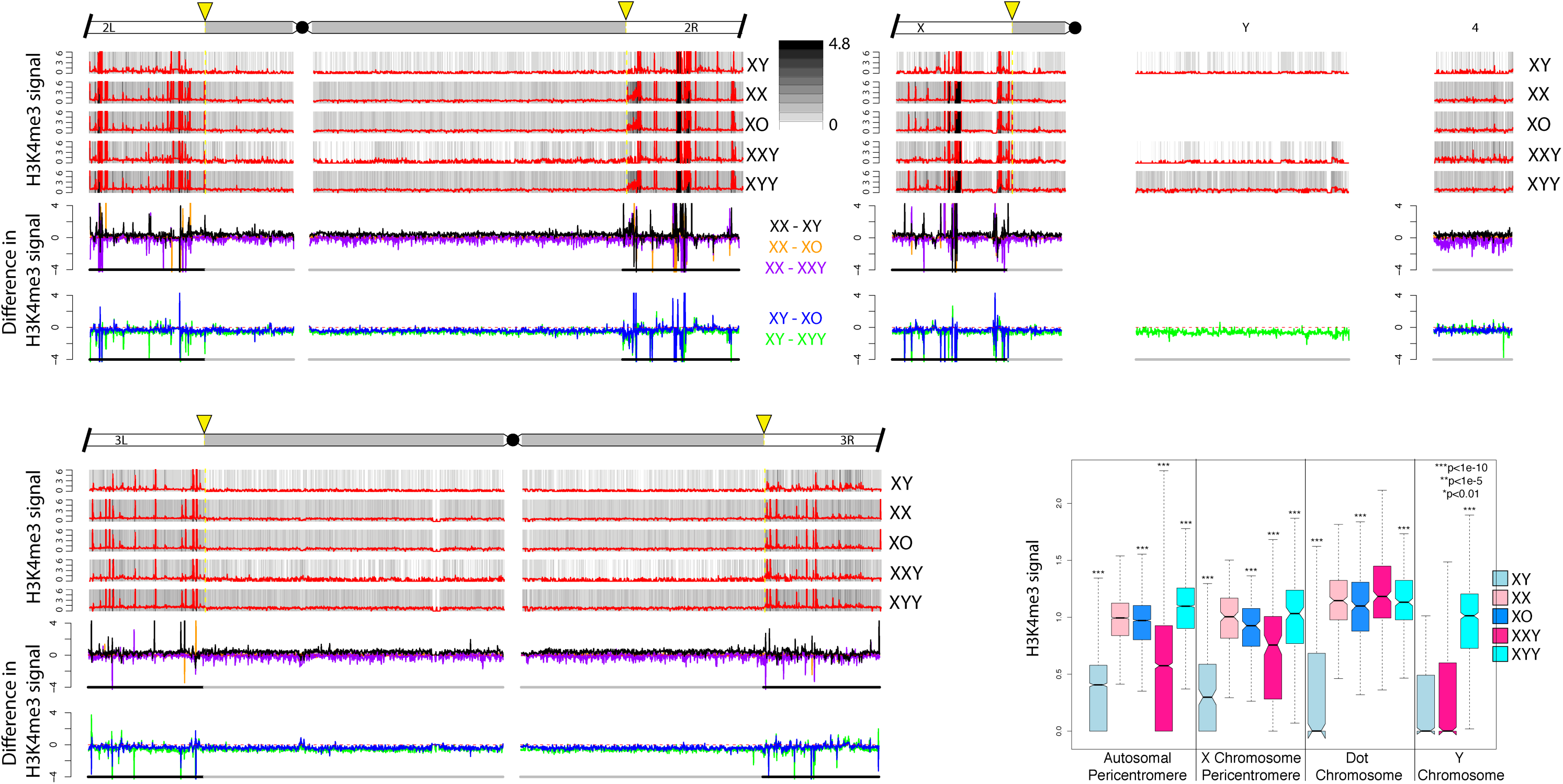
Enrichment of H3K4me3 for *D. melanogaster* strains with different karyotypes across the euchroma3n / heterochroma3n boundary along each chromosome arm. We show the centromere-proximal 1Mb euchroma3c region of chromosome 2, 3 and X, as well as the complete assembled heterochroma3n region for each chromosome. For each karyotype, the enrichment in 5kb windows is shown in red lines (normalize ra3o of ChIP to input, see Materials & Methods), and the same data is shown in gray scale according to the scale in the upper right. Note that the enrichment profiles for all 5 karyotypes are ploeed on the same scale to allow for direct comparisons. Below the enrichment profiles for each chromosome arm, subtrac3on plots show the absolute difference in signal of 5kb windows between pairs of karyotypes along the chromosome arms. The cytogenomically defined heterochroma3n is marked by gray bars, the the euchroma3n / heterochroma3n boundary is indicated by a yellow arrow. The box plots show the ChIP signal for all 5kb windows in different chromosomal regions, with boxes extending from the first to the third quar3le and whiskers to the most extreme data point within 1.5 3mes the interquan3le range. P-values were calculated rela3ve to XX females for XY males and XXY females, and rela3ve to XY males for XO and XYY males; p-values for the Y chromosome were calculated rela3ve to XY males (Wilcoxon test). For genome-wide enrichment plots see **Fig S2**.

**Figure 3.**
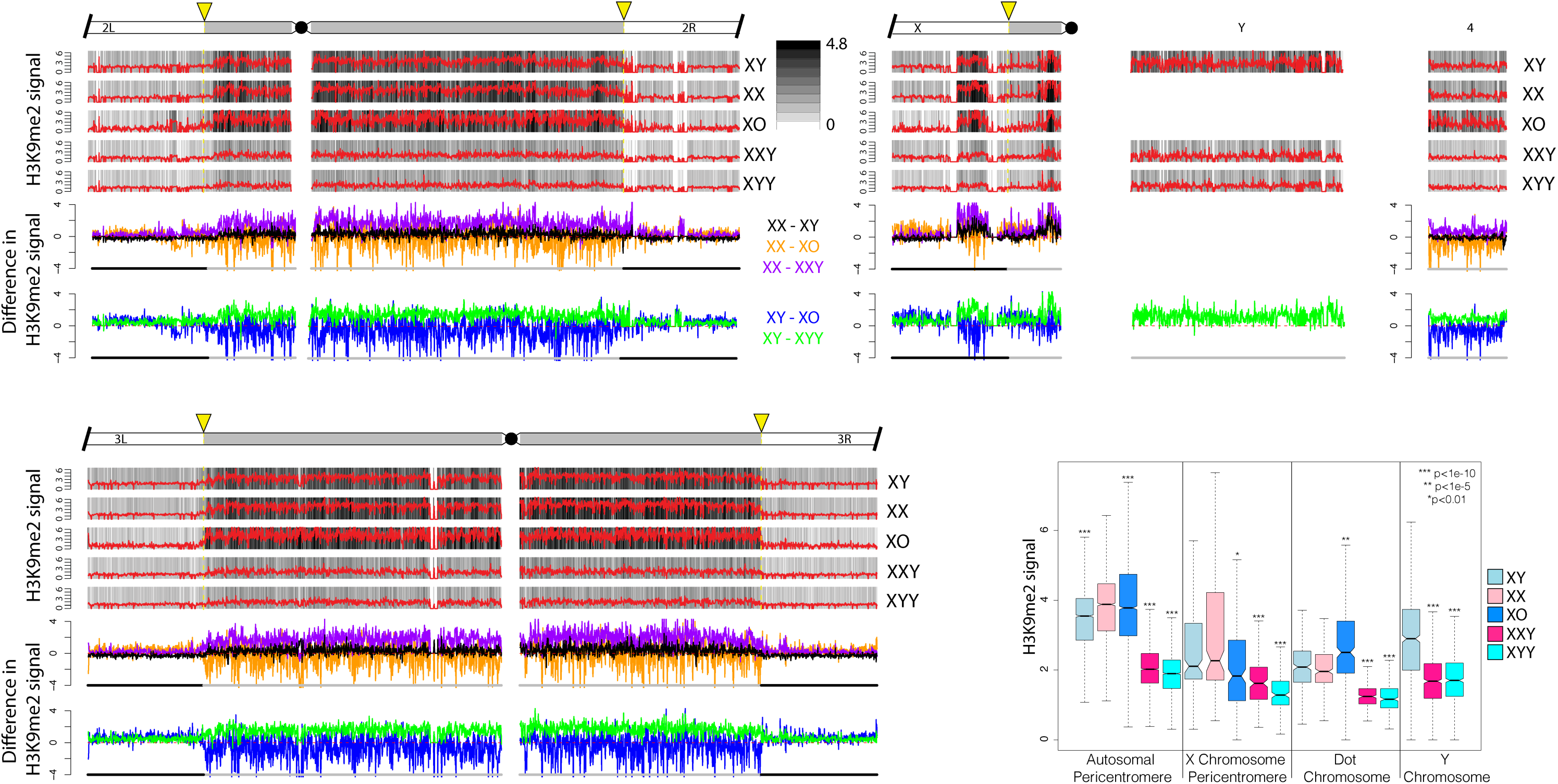
Enrichment of H3K9me2 for *D. melanogaster* strains with different karyotypes across the euchroma3n / heterochroma3n boundary along each chromosome arm. We show the centromere-proximal 1Mb euchroma3c region of chromosome 2, 3 and X, as well as the complete assembled heterochroma3n region for each chromosome. For each karyotype, the enrichment in 5kb windows is shown in red lines (normalize ra3o of ChIP to input, see Materials & Methods), and the same data is shown in gray scale according to the scale in the upper right. Note that the enrichment profiles for all 5 karyotypes are ploeed on the same scale to allow for direct comparisons. Below the enrichment profiles for each chromosome arm, subtrac3on plots show the absolute difference in signal of 5kb windows between pairs of karyotypes along the chromosome arms. The cytogenomically defined heterochroma3n is marked by gray bars, the euchroma3n / heterochroma3n boundary is indicated by a yellow arrow. The box plots show the ChIP signal for all 5kb windows in different chromosomal regions, with boxes extending from the first to the third quar3le and whiskers to the most extreme data point within 1.5 3mes the interquan3le range. P-values were calculated rela3ve to XX females for XY males and XXY females, and rela3ve to XY males for XO and XYY males; p-values for the Y chromosome were calculated rela3ve to XY males (Wilcoxon test).

**Figure 4.**
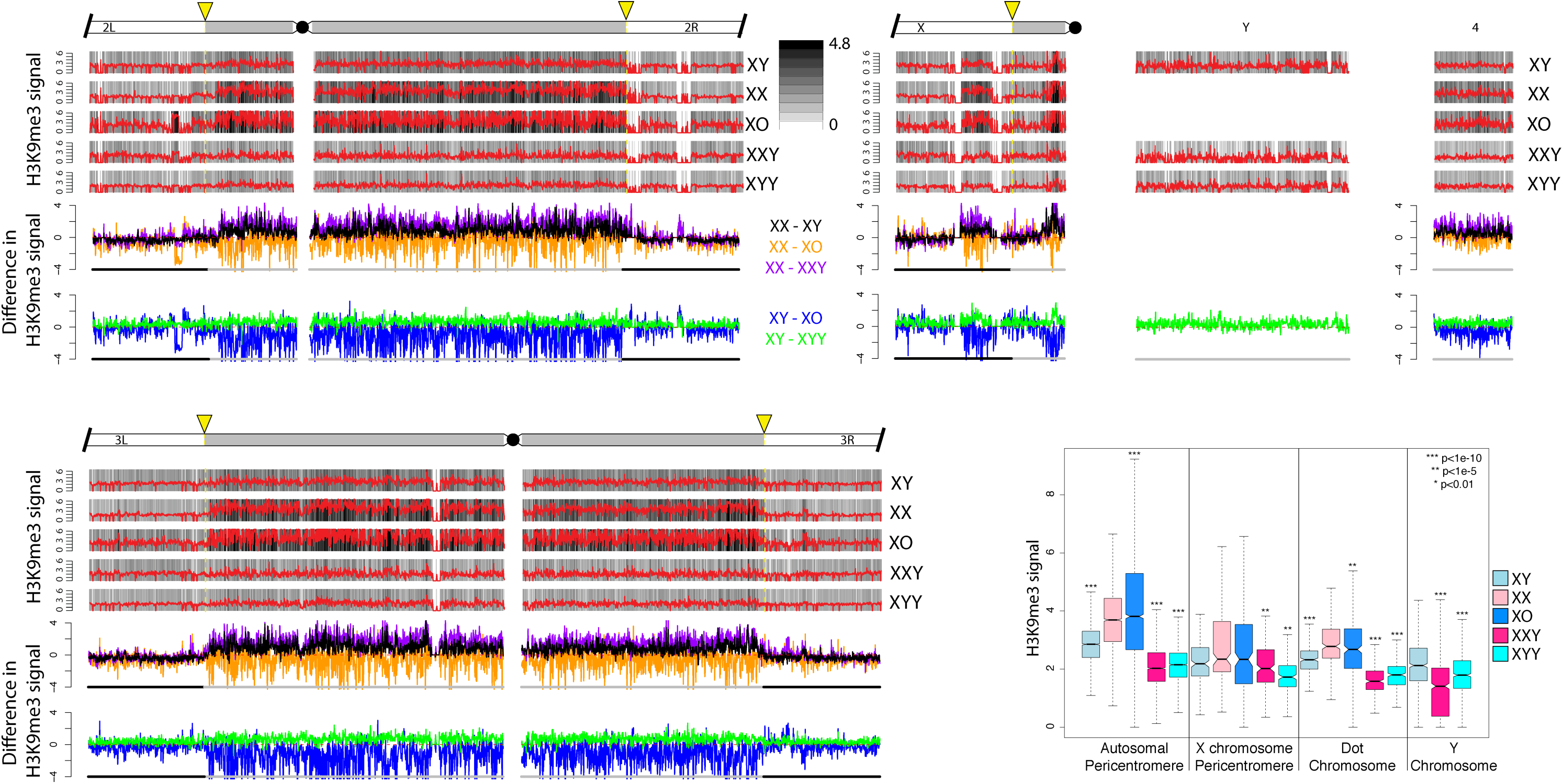
Enrichment of H3K9me3 for *D. melanogaster* strains with different karyotypes across the euchroma3n / heterochroma3n boundary along each chromosome arm. These plots were made in the same manner as those for H3K9me2 (see Figure 2).

### The genomic distribution of active chromatin is similar in flies with different karyotypes

The histone modification H3K4me3 primarily associates with active genes (Guenther, et al. 2007; Kharchenko, et al. 2011) and is highly underrepresented in repeat-rich regions, including the Y chromosome; we thus expect that its relative abundance and genomic distribution is little influenced by the dose of Y chromosomes. Indeed, we find that H3K4me3 peaks are primarily located along the euchromatic chromosome arms, and highly deficient in pericentromeric regions, and along the Y chromosome (**Figure S3;** for zoom-in at the heterochromatin / euchromatin boundary see **Figure 2**). Genomic enrichment patterns of H3K4me3 are similar across sexes and flies with varying numbers of Y chromosomes (**Figure 2, S3**), both when comparing the relative position of peaks, but also the absolute magnitude of signal across samples (**Figure 2, S3**). This confirms our expectation that Y dose should not dramatically influence the distribution of active chromatin marks, and also suggests that our normalization procedure is accurate in quantifying relative abundance of histone modifications across samples. Western blots confirm our inferences based on ChIP-seq, i.e. that H3K4me3 signal is similar across flies with different karyotypes (**Figure S4**).

### Heterochromatic histone modifications in wildtype flies

We investigated the genomic distribution of two histone marks that are associated with heterochromatin formation, H3K9me2 and H3K9me3 (Kharchenko, et al. 2011). If the Y chromosome indeed acts as a sink for components of the heterochromatin machinery, we expect global differences in the enrichment patterns of heterochromatic histone modifications across strains with different numbers of Y chromosomes, or more generally, across flies with different amounts of repetitive DNA (see Figure 1A, B). Specifically, we expect that as the repeat content increases, flies should harbor less H3K9me2/3 at their heterochromatic regions. In wildtype Drosophila, heterochromatin is highly enriched in pericentromeric regions, the small dot chromosome, and along the entire length of the Y chromosome (Bonaccorsi and Lohe 1991; Gatti and Pimpinelli 1983; Hannah 1951). Note that the *D. melanogaster* Y chromosome is estimated to be about 40Mb (i.e. 20% of the haploid male genome; Gatti and Pimpinelli 1983; Hoskins, et al. 2002), but only 3.7Mb (i.e. <10%) of the Y chromosome has been assembled. Similarly, other heterochromatic regions are also only partly assembled: 1.5Mb (∼25% of the pericentromeric heterochromatin) on chromosome 2L, 5.4Mb (∼50%) on 2R, 5.1Mb (∼50%) on 3L, 4.2Mb (∼50%) on 3R and only 0.9Mb (i.e. only about 5% of the pericentromeric heterochromatin) on the X chromosome. Thus, our genome mapping analysis will underestimate the extent of heterochromatic histone modifications that are associated with the Y chromosome, and other repetitive regions. Also note that the pericentromeric heterochromatin along the X chromosome is non-continuous (Figs. 2, 3; see also Riddle, et al. 2011), with the more distal heterochromatic block encompassing the *flamenco* locus (Goriaux, et al. 2014).

Overall, we find that levels of heterochromatin enrichment are similar for the H3K9me2 and H3K9me3 marks, but differ between flies with varying amount of repetitive DNA (Figure 3, 4; for genome-wide plots see **Figures S5**). The male-specific Y chromosome is highly enriched for both of these repressive histone modifications in wildtype males, and we find that wildtype females have slightly higher levels of H3K9me2/3 enrichment than males in their pericentromeric regions, and on the dot chromosome, relative to euchromatic background levels (Figure 3, 4). Moreover, the heterochromatin / euchromatin boundary is slightly less clearly discernable from H3K9me2/3 enrichment patterns for males relative to females (**Figure 5, Figure S6, S7**). Western blots suggest that males harbor slightly more H3K9me2/3 compared to females (**Figure S4**). Thus, we find strong enrichment of the heterochromatic histone modifications on the Y and their relative deficiency at pericentromeric regions on autosomes and the X in wildtype males relative to females, despite similar amounts of overall H3K9me2/3. This observation is consistent with the hypothesis that the repeat-rich Y chromosome acts as a sink for components of the heterochromatic machinery, resulting in a relative paucity of heterochromatic histone modifications elsewhere in the genome. However, despite quantitative differences in levels of heterochromatic histone modifications, overall patterns of H3K9me2/3 enrichment are similar between sexes.

**Figure 5.**
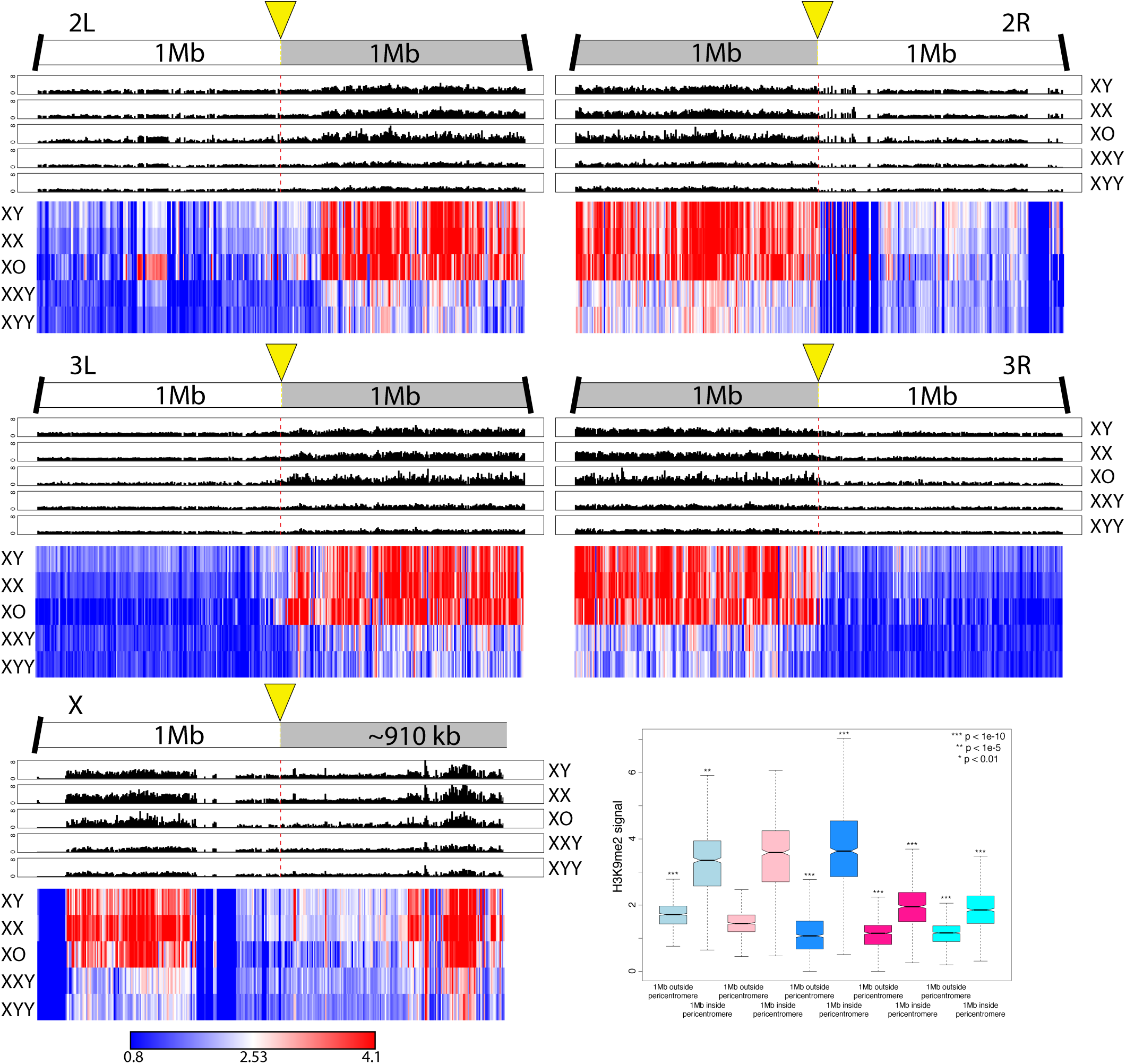
Enrichment of H3K9me2 within 1Mb of the heterochroma3n/ euchroma3n boundaries (as defined in the Release 6 of the *D. melanogaster* genome (Hoskins, et al. 2015)). The upper panels show H3K9me2 signal in 5kb windows for each chromosome arm, and the boeom panel shows scaled heatmaps for the same 5kb windows, to allow direct comparisons of H3K9me2 signal across samples. For H3K9me3 plots, see **Figure S3**. Box plots show H3K9me2 signal of 5kb windows in euchromatic regions 500kb outside the pericentromere vs. 500kb inside the heterochroma3n/ euchroma3n boundary. Significance values are all calculated using the Wilcoxon test.

### Heterochromatic histone modifications in X0, XXY & XYY flies

To investigate the Y chromosome’s role in the genome-wide distribution and enrichment for heterochromatic components, we studied histone modification profiles from female flies containing a Y chromosome (XXY females), and males with either zero or two Y chromosomes (X0 vs. XYY males). Female Drosophila that contain a wildtype Y chromosome show clear enrichment for both heterochromatic histone modifications on the Y chromosome, but an overall reduction in levels of H3K9me2/3 relative to wildtype females, both at pericentromeric regions and along the dot (Figure 3, 4). The genomic distribution of H3K9me2/3 in XXY females is consistent with the model of the Y chromosome acting as a sink for components of the heterochromatin machinery, sequestering heterochromatic proteins to the Y chromosome and diluting them away from autosomal and X-linked targets. XXY females also show less heterochromatic histone modifications at pericentromeric regions and the dot relative to wildtype XY males (Figure 3, 4). This is consistent with the higher repeat content in XXY flies compared to XY flies -due to the large heterochromatic block on the X -contributing to the heterochromatin sink effect. This suggests that the effect of the Y chromosome on heterochromatin distribution is not a unique property of the Y but instead a result of a large amount of any additional repetitive sequence. XYY males harbor the highest amount of repetitive DNA and show severely decreased levels of H3K9me2/3 enrichment along repeat-rich, normally heterochromatic regions, including their Y chromosomes, pericentromeric regions, and along the dot, relative to levels found in other karyotypes investigated (Figure 3, 4).

X0 males, on the other hand, have the lowest repeat content of all flies, and show the strongest enrichment of heterochromatic histone modifications at pericentromeric regions and along the dot chromosome (Figure 3, 4). Enrichment levels of H3K9me2/3 at repetitive regions (pericentromere and the dot) relative to euchromatic background levels in X0 males is well above that of wildtype males and also wildtype females (or XXY females, which have the same autosomal background as X0 flies; Figure 3, 4). Similar patterns of re-distribution of heterochromatin are observed in a biological replicate without a spike (**Figure S8**), and in X0, XXY, and XYY flies that were generated using a different attached X stock, 4248 (**Figure S9**), demonstrating that our findings are robust in different genetic backgrounds. Together, our data provide clear evidence that Y chromosomes, and repetitive DNA in general, affect heterochromatin formation genome-wide, consistent with a model of the Y chromosome or other large blocks of repetitive sequences acting as heterochromatin sinks, possibly by redistributing heterochromatin components across the genome. Note that the sink effect of additional heterochromatin is almost linearly related to increasing amounts of repetitive DNA: We see increasingly less heterochromatin form at the pericentromeres and the dot chromosome as the repeat content increases, with X0 < XX < XY < XXY < XYY (see boxplot in Figure 3, 4).

The depletion of heterochromatic histone modifications from pericentromeric regions causes the euchromatin/ heterochromatin boundaries to be significantly diluted in XXY and XYY individuals (**Figure 5**, **Figure S6, S7**). X0 males, in contrast, show spreading of their pericentromeric heterochromatin into chromosome arms that are normally euchromatic in wildtype flies, which is consistent with previous studies that found enhanced position effect variegation in XO males (**Figure 5, Figure S6, S7;** Belyaeva, et al. 1993; Wallrath and Elgin 1995b).

Overall, we see that increasing the amount of repetitive DNA by changing the dose of both sex chromosomes corresponds with a decrease in the signal of heterochromatic histone modifications at pericentromeric regions and along the dot chromosome. This is consistent with a model of stoichiometric balance between protein components involved in the formation of heterochromatin and the amount of repetitive DNA sequences within a genome. Together, ChIP-seq profiles of histone modifications in wildtype flies, X0 and XXY males, and XXY females support the hypothesis that the Y chromosome acts as a heterochromatin sink in Drosophila.

### Sex chromosome dose and gene expression

Polymorphic Y chromosomes affect expression of hundreds of autosomal and X-linked genes in *D. melanogaster*, a phenomenon known as Y-linked regulatory variation (YRV) (Dimitri and Pisano 1989; Lemos, et al. 2008; Lemos, et al. 2010; Sackton, et al. 2011). To test if genes that respond to YRV are also expressed differentially in flies with different sex chromosome configurations, we collected biologically independent replicate RNA-seq data from heads for wildtype males and females, as well as from X0, XXY and XYY flies. As noted above, protein-coding Y-linked genes in Drosophila are only expressed in male germ line and thus cannot directly contribute to differences in expression profiles in head samples among flies with different numbers of Y chromosomes. Overall, we find that 100s of genes show differential expression among flies with different sex chromosome karyotypes (**Figure 6A**). GO analysis revealed that differentially expressed genes tend to be enriched for functions associated with reproductive processes (**Table S4**), and are not simply clustered around pericentromeric regions (**Figure S10, S11**). Genes that are expressed most differently between XO and XY males, and XX and XXY females, show significantly greater difference in H3K9me2 signal compared to all genes, while these genes have significantly less difference in H3K4me3 signal compared to all genes (**Figure S12**). This is consistent with the hypothesis that the Y chromosome re-distributes heterochromatin components, and can thereby influence the expression of hundreds of genes. However, we see no global relationship between gene expression differences and H3K9me2/3 enrichment levels across all genes (**Figure S13**). This suggests that the effect of the Y chromosome on heterochromatin and its effect on gene expression are not explained by a simple model whereby the Y chromosome modifies heterochromatin formation and thereby directly modifies gene expression across the entire genome. Indeed, the majority of genes are not targeted by H3K9me2/3 above background levels in any of the karyotypes investigated, and thus those marks are unlikely to directly influence global gene expression patterns.

**Figure 6.**
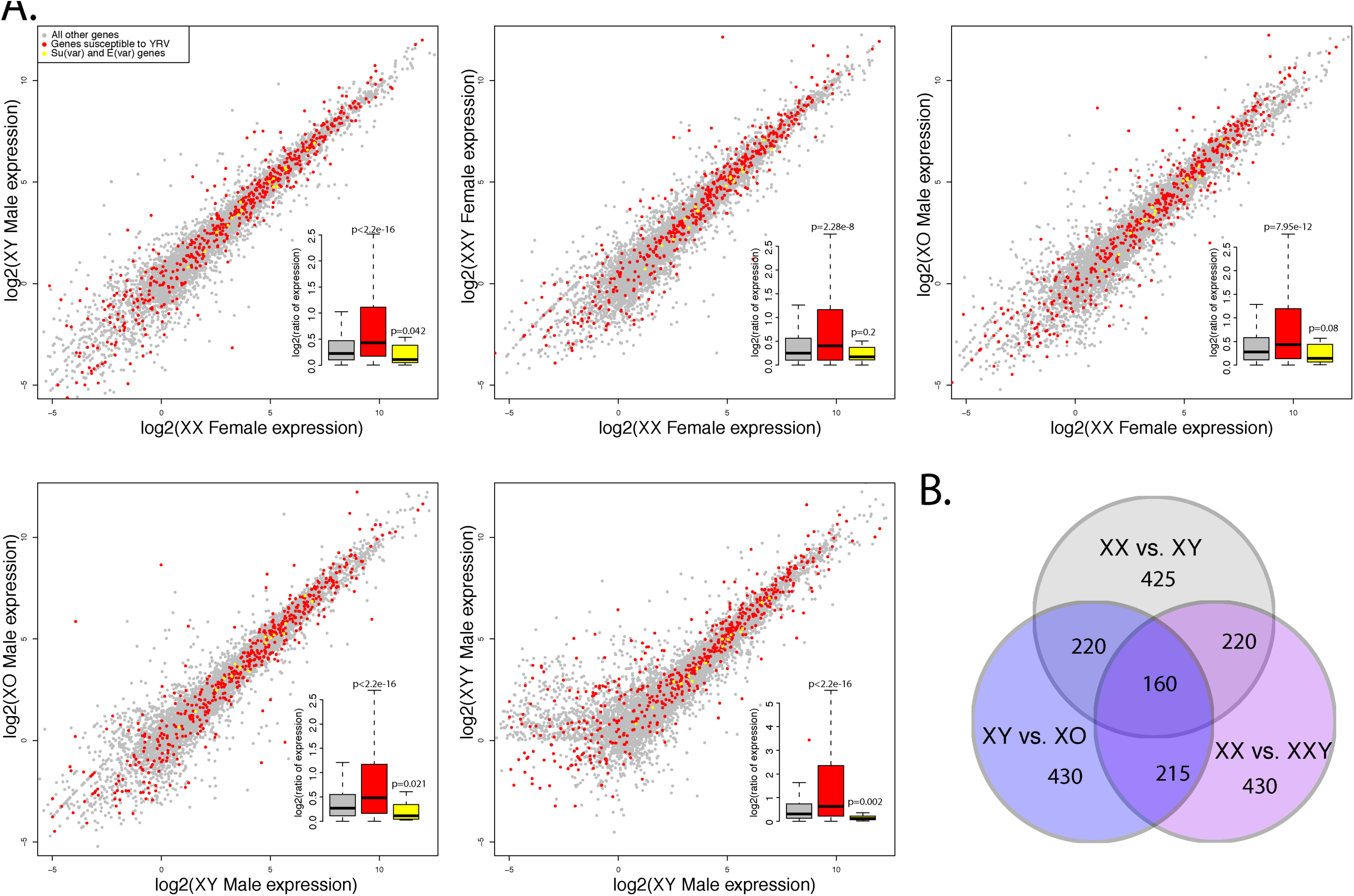
Gene expression varia3on between flies with different sex chromosome karyotypes. **A.** Pairwise expression comparisons for flies with different karyotypes. Genes marked in red are susceptible to YRV (from (Sackton and Hartl 2013)), and genes in yellow are genetically defined Su(var) and E(var) genes in *D. melanogaster* (from (Elgin and Reuter 2013)); gray genes are all other genes. **B.** Overlap of top 10% of differentially expressed genes between wildtype XY male and XX female, and males and females with and without Y chromosomes, i.e. XX vs. XXY females and XY vs. X0 males

We used a consensus set of 678 genes that were classified as susceptible to YRV (Sackton and Hartl 2013), and found that these genes were generally expressed more differently between different sex chromosome karyotypes compared to random genes (**Figure 6A**). This suggests that a similar mechanism is underlying both YRV and gene expression differences in flies with different sex chromosome configurations. Genes that are genetically defined to either suppress or enhance silencing in assays for PEV in *D. melanogaster*, i.e. Su(var) and E(var) genes (Elgin and Reuter 2013), are expressed at similar levels in flies with different karyotypes (**Figure 6A**). This is consistent with our Western blots that reveal no consistent differences in H3K9me2/3 levels among flies with different sex chromosome configurations (**Figure S3**).

Interestingly, genes susceptible to YRV are more likely to be differentially expressed between wildtype sexes, and genes that are differentially expressed between males and females in head tissue tend to also be differentially expressed between X0 and XY males, or XX and XXY females (p<1e-6, permutation test, **Figure 6B, Figure S14**). In particular, 160 of the top 10% genes that are differentially expressed between wildtype XX females and XY males, vs. X0 and XY males vs. XX and XXY females overlap, while we only expect 9 by chance. This suggests that a substantial fraction of sex-biased expression in somatic tissues may simply be an indirect consequence of the absence or presence of the Y, i.e. the sink effect of the Y chromosome may contribute to sex-biased expression patterns in *D. melanogaster*.

### Repeat reactivation in XXY and XYY flies

Heterochromatin is established during early embryogenesis and leads to the transcriptional silencing of repetitive DNA and transposable elements (TEs) (Elgin and Reuter 2013). We used our RNA-seq data to assess whether changes in chromatin structure due to Y chromosome dose are associated with changes in gene expression patterns of repetitive elements. We first used consensus sequences of known TEs annotated by FlyBase (flybase.org), and found that overall repeat content correlated negatively with H3K9me2/3 enrichment at TEs: X0 flies had the highest level of H3K9me2/3 enrichment across TE families, followed by XX and XY wildtype flies, and XXY and XYY flies having the lowest amount of heterochromatin marks at their TEs (p<0.01 for each comparison; **Figure 7A, Figure S15A**; note that these estimates are corrected for differences in copy numbers between repeats, by looking at the enrichment of H3K9me2/3 enrichment over input for each karyotype). Despite dramatic differences in overall levels of repressive histone marks across repeat families, levels of expression for the various TEs between karyotypes are very similar (p>0.05, **Figure 7A, Figure S16**). A subset of TEs shows an increase in expression in XYY males compared to other samples, including at least 5 retroviral elements (1731, 297, Max element, mdg1, and mdg3, **Figure S16**). Increased expression of these repeats appears in part be driven by an increased copy number in the XYY male genome; if we correct for genomic copy number, we find that only three of these repeats (1731, 297, and Max element) are expressed more highly in XYY males compared to the other karyotypes (**Figure S17**). Thus, despite global differences in heterochromatin formation associated with repeats across karyotypes, this does not manifest itself in a global de-repression of TEs, but seems to instead involve de-repression of just a subset of TE families. Note that a loss of heterochromatin at pericentromeric regions and the Y chromosome should not necessarily result in increased expression across all TE families present. On one hand, most TEs located in the pericentromere and the Y chromosome are partial and non-functional copies that have lost their ability to transpose (Ananiev, et al. 1984; Pimpinelli, et al. 1995). In addition, we assayed gene expression in somatic head tissue, and many TE families only mobilize in the germline (Charlesworth and Langley 1989).

**Figure 7.**
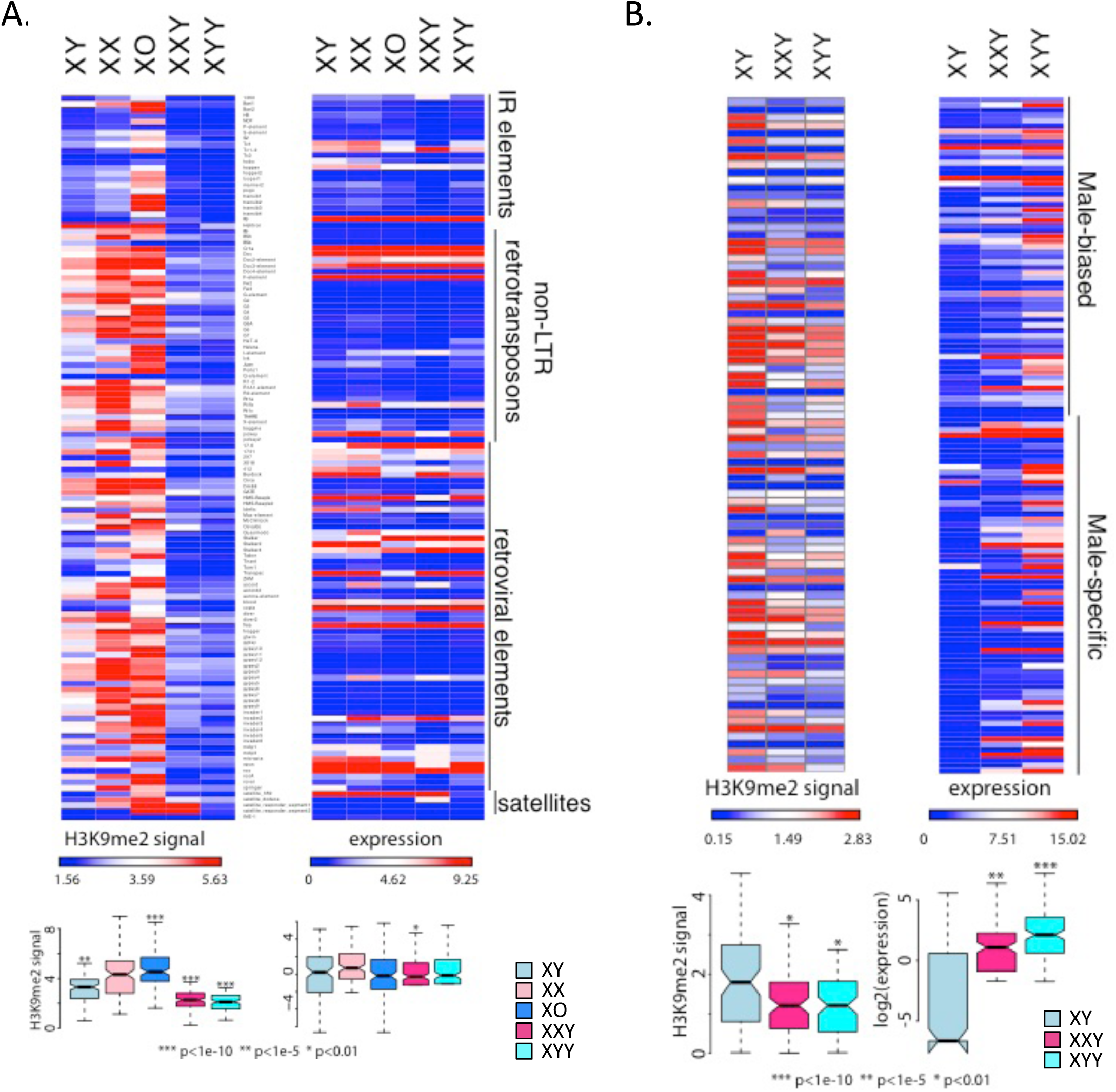
Chroma3n and expression paeerns at TE families. Shown is enrichment of H3K9me2 at different TE families (rela3ve to genome-wide levels) and their expression levels for the various karyotypes. For H3K9me3 plots, see **Figure S8**. **A.** All repeats from the library of consensus transposable elements and satellites from FlyBase. Boxplots summarize H3K9me2 and expression values across all repeats, and significance values were calculated using the Wilcoxon test; all comparisons of expression levels were not significant. **B.** Puta3vely Y-linked (male-specific) *de novo* assembled repeats only. Boxplots summarize H3K9me2 and expression values across all puta3vely Y-linked repeats, and p-values were calculated rela3ve to XY males using the Wilcoxon test.

Most of the Y chromosome has not yet been assembled (Hoskins, et al. 2015), including its repetitive elements, and we were interested in whether expression of Y-linked repeats would be particularly sensitive to Y chromosome dosage. We thus used a *de novo* approach to identify male-specific Y-linked repeats that does not rely on a genome assembly, but instead uses kmer abundances from next generation sequencing reads to produce a repeat library (Koch, et al. 2014). We then mapped male and female genomic reads from the Canton-S strain back to our *de novo* assembled repeat library, in order to infer Y-linkage for repeats that were only covered by male genomic reads (**Figure S18, S19**). Male-specific repeats are highly enriched for H3K9me2/3 in wildtype males, and transcriptionally silenced (**Figure 7B**). However, while Y-linked repeats show similar enrichment for the H3K9me3 mark in all karyotypes (**Figure S15B**), XXY females and XYY males are highly deficient for H3K9me2 at Y-linked repeats and expression of Y-linked repeats is de-repressed relative to wildtype males (**Figure 7B, Figure S20**). If we account for differences in copy number of the Y-linked repeats, we still find that Y-linked repeats are expressed more highly in XXY females and XYY males compared to wildtype males (**Figure S21**). Thus, consistent with the ChIP-seq data that showed low levels of heterochromatic histone modifications (especially H3K9me2) along the Y of XXY females or the two Y chromosomes of XYY males, relative to wildtype males, our gene expression data demonstrate that Y-linked repeats become transcriptionally activated in female flies that normally do not have a Y chromosome, or male flies with double the dose of Y-linked repeats, and this is not simply a consequence of an increased copy number of Y-linked repeats.

## Discussion

### Dosage effects of chromatin components and repetitive DNA

Many eukaryotic genomes contain large amounts of selfish, repetitive DNA; in the Drosophila strains investigated, for example, the heterochromatin content varies from about ⅓ to ½ of the genome (see **Fig. 1B**). Transcriptional silencing of repeats through heterochromatin formation is one way to alleviate the deleterious effects of repetitive DNA (Elgin and Reuter 2013). Studies of PEV in *D. melanogaster* have yielded important insights into the biology of heterochromatin (Muller 1930; Schultz 1936; Zhimulev, et al. 1986), and frequently found dose-dependent effects of chromatin proteins and trans-activating factors (Weiler and Wakimoto 1995). For example, depletion of HP1, an important protein involved in both the recruitment and maintenance of heterochromatic histone modifications, suppresses variegation (i.e. it results in less heterochromatin formation and thus less suppression at a reporter gene; Eissenberg, et al. 1990), whereas increased dosage of HP1 enhances variegation (i.e. it increases silencing through increased heterochromatin formation; Eissenberg, et al. 1992). Addition of extra heterochromatin to the genome suppresses variegation, while its subtraction enhances variegated gene expression (Baker 1968; Gatti and Pimpinelli 1992; Spofford 1976). In *D. melanogaster*, the Y chromosome is a potent suppressor of variegation, i.e. it induces less heterochromatin at a reporter gene (Gowen and Gay 1934), and *D. melanogaster* males with different Y chromosomes in otherwise identical genetic backgrounds vary in their propensity to silence a heterochromatin-sensitive reporter gene in PEV assays (Lemos, et al. 2010). Highly repetitive Y chromosomes are thought to sequester heterochromatic factors that are present in only limited amounts (Dimitri and Pisano 1989), and different Y chromosomes vary in their repeat content and thus the extent to which they sequester those heterochromatin components, thereby influencing PEV. Reporter gene assays, however, do not directly probe the integrity or amount of heterochromatin at repetitive regions that exert PEV through spreading of heterochromatin, nor do they directly demonstrate changes in the spreading of heterochromatin along the chromosome (Dimitri and Pisano 1989; Gatti and Pimpinelli 1992; Spofford 1976). Under the heterochromatin sink model, changes in the amount of repetitive DNA should globally modify the amount of heterochromatin formed at repeats. Specifically, decreasing the amount of repetitive DNA should result in increased levels of heterochromatin at repetitive regions, since heterochromatic factors that are present in only limited amounts can be sequestered at the existing repeats at higher concentration, while increased amounts of repeats would result in the opposite effect.

In our study, we directly demonstrate that the Y chromosome, and repeat-rich DNA in general, can act to globally affect heterochromatin formation in *D. melanogaster*. Consistent with the heterochromatin sink model, we find that increasing the amount of repetitive DNA generally decreases the amount of H3K9me2/3 enrichment at repeat-rich regions, such as pericentromeres, the dot, or the Y chromosome. Individuals with the lowest repeat content (X0 males in our experiment) show the highest enrichment of H3K9me2/3 in repeat-rich regions, and the pericentromeric heterochromatin on the autosomes of X0 flies clearly extends into genomic regions that are normally euchromatic in wildtype *D. melanogaster*. Wildtype females show slightly higher H3K9me2/3 levels at their pericentromeric regions and the dot chromosome and a slightly sharper euchromatin/ heterochromatin boundary at autosomes compared to wildtype males. Indeed, females generally show a higher degree of silencing in assays for PEV, suggesting that normally euchromatic regions are more prone to acquire a heterochromatic conformation in females (Girton and Johansen 2008; Wallrath and Elgin 1995a).

XYY males and XXY females, on the other hand, show a dramatic reduction of H3K9me2/3 enrichment at repeat-rich regions, and the boundaries between the heterochromatic pericentromere and the euchromatic chromosome arms become blurry. Overall, the sink effect of additional heterochromatin appears almost linearly related to increasing amounts of repetitive DNA, consistent with studies based on PEV (Dimitri and Pisano 1989). Thus, this dosage sensitivity of H3K9me2/3 enrichment in repetitive regions suggests that there is a stoichiometric balance among protein components and total repeat content of the genome to maintain proper heterochromatic silencing.

### Functional heterogeneity of heterochromatin

Most DNA sequences that comprise the various heterochromatic elements are not unique and specific to chromosomes of chromosomal segments, but are shared with other genomic regions; most satellite DNA repeats map to multiple genomic sites (Bonaccorsi and Lohe 1991), and so do nearly all the transposable elements (Pimpinelli, et al. 1995). All highly repetitive blocks in the *D. melanogaster* genome, including both arms of the Y chromosome, the heterochromatic segments located at the base of the X chromosome, the left and right arms of chromosomes 2 and 3 and the fourth chromosome heterochromatin are all effective in inducing PEV (Spofford 1976). An important open question is whether the ability to suppress variegation is a general property of all heterochromatic regions, or whether it can be ascribed to specific heterochromatic sites. PEV suppression exerted by the Y chromosome was mapped using a variety of cytologically determined Y-chromosomal deficiencies and Y-linked fragments (Dimitri and Pisano 1989). The suppression effect exerted by the Y chromosome was found to be positively related to the size of the Y-derived DNA, and was not attributable to any discrete Y region; Y chromosomes that were cytologically different yet retain similar amounts of heterochromatin were found to be equally effective suppressors (Dimitri and Pisano 1989). Thus, at the level of resolution provided by these cytogenetic studies, all Y-fragments appear to be similarly effective in influencing global chromatin structure, as assayed by PEV assays. In our study, we also find that there is a clear inverse relationship between the amount of repetitive DNA present, and the amount of heterochromatin induced at repeats, by varying the repeats derived from both the X and the Y chromosome. This is consistent with the notion that there is limited heterogeneity among repeats in influencing global chromatin structure. However, it will be of great interest to more carefully characterize the effects of specific repeat elements on the Y chromosome, such as the rDNA cluster, or specific types of satellites, to directly address the question of how uniform the sink effect is across different repeat types.

### Functional consequences of the Y chromosome’s global effects on heterochromatin

Analyses of gene expression profiles suggest that global changes in heterochromatic histone modifications can have broad functional consequences for the organism. Specifically, we show that hundreds of genes are differentially expressed in individuals that differ in their sex chromosome karyotype, and genes that are susceptible to YRV are more prone to be differentially expressed in individuals with different sex chromosome complements. We find that increasing the amount of repetitive DNA leads to a decrease in heterochromatic histone modification signal at TEs. XYY males and XXY females have low levels of H3K9me2 signal in TEs, and especially so in male-specific repeats, and we show that this deficiency of heterochromatin is associated with a de-repression of Y-linked repeats that we detect as an increase in expression levels of these repeats. Thus, while fruit flies have efficient mechanisms in place to silence wildtype levels of repetitive DNA, a large increase in the amount of repetitive sequences, caused by introducing additional Y chromosomes, limits the organism’s ability to form heterochromatin and those additional repeats apparently cannot be efficiently silenced.

Whole-genome sequencing studies can provide information on the genome size by estimating the amount of euchromatic DNA, but cannot reliably estimate the amount of repetitive, heterochromatic sequences. Cytogenetic studies suggest that individuals within a population can differ greatly in how much repetitive heterochromatic DNA they contain. The size of the pericentromeric heterochromatic block on the *D. melanogaster* X chromosome, for example, varies by about 2-fold among strains (Halfer 1981), and dramatic variation in size and morphology of the Y chromosome has been reported in natural populations of *D. pseudoobscura* (Dobzhansky 1937). Moreover, haploid genome size estimates of different *D. melanogaster* strains using flow cytometry differ by almost 100Mb, and the vast majority of this variation is thought to result from differences in repetitive heterochromatin (Bosco, et al. 2007). Similarly, a recent bioinformatics analysis that identified and quantified simple sequence repeats from whole genome sequences also found a 2.5-fold difference in their abundance between *D. melanogaster* strains (Wei, et al. 2014). Thus, natural variation in repetitive DNA among individuals may in fact span a wider range than that across sex chromosome karyotypes investigated here. This implies that repetitive DNA might serve as an important determinant of global chromatin dynamics in natural populations, and may be an important modifier of the differential expression of genes and TEs between individuals.

### Heterochromatin/ euchromatin balance between sexes

Males contain a Y chromosome that is highly repetitive and heterochromatic, and which may shift the genome-wide heterochromatin/ euchromatin balance between the sexes (Brown and Bachtrog 2014). In particular, if the Y chromosome sequesters proteins required for heterochromatin formation, males may be more sensitive to perturbations of the balance between repetitive sequence content and heterochromatic protein components, and might have lower levels of heterochromatin-like features in the rest of their genome, as compared to females (Brown and Bachtrog 2014). Indeed, RNAi knockdown of the heterochromatin protein HP1 preferentially reduces male viability (Liu, et al. 2005), and the presence of Y-linked heterochromatin is thought to underlie this differential sensitivity.

Heterochromatin formation is temperature-sensitive, and female Drosophila are more tolerant of heat shock, survive heat-induced knock-down better (Yamamoto and Ohba 1982), and become sterile at higher temperatures than males (David, et al. 2005), and it is possible that differences in the chromatin landscape may contribute to sex-specific differences in heat stress response. Indeed, the Y chromosome was found to be responsible for much of the genetic variation of heat-induced male sterility found across populations (Rohmer, et al. 2004). Also, as mentioned, female flies show stronger silencing in assays for PEV (Girton and Johansen 2008; Wallrath and Elgin 1995a), consistent with having more heterochromatin protein components relative to repetitive sequences, which can then spread into reporter genes more readily.

Many recent studies in animals have shown that a large portion of the transcriptome in animals is sex-biased (Mank, et al. 2008; Ranz, et al. 2003). Sex-biased expression patterns are typically seen as an adaptation to form the basis of sexually dimorphic phenotypes (Parsch and Ellegren 2013). In Drosophila, most sex-biased expression patterns are due to differences in expression in sex-specific tissues (i.e. gonads; Assis, et al. 2012; Meisel 2011); however, hundreds of genes also show differential expression in shared, somatic tissues (Assis, et al. 2012; Meisel 2011). Interestingly, we find that a similar set of genes that show differences in expression patterns between males and females (in head) are also differentially expressed between XY and X0 males, or XX and XXY females. This suggests that not sex *per se*, but the absence or presence of the Y chromosome is responsible for much of the differences in expression patterns between sexes. Thus, sex-biased gene expression is normally interpreted as a sex-specific adaptation to optimize expression levels of genes in males and females (Parsch and Ellegren 2013). However, our results suggests that it is also possible that sex-biased expression patterns are simply an indirect consequence of global differences in the chromatin structure between males and females, due to the presence of a large repetitive Y chromosome in males.

## Materials & Methods

### Drosophila strains

Fly strains were obtained from the Bloomington Stock Center. The following strains were used: Canton-S, 2549 (C(1;Y),y^1^cv^1^v^1^B/0 & C(1)RM,y^1^v^1^/0), and 4248 (C(1)RM,y^1^pn^1^v^1^/C(1;Y),y^1^B^1^, y^1^B^1^/0;sv^spa-pol^).. The crossing scheme used to obtain X0 and XYY males and XXY females is depicted in **Figure 1B**. For chromatin and gene expression analyses, flies were grown in incubators at 25°C, 32% of relative humidity, and 12h light. Newly emerged adults were collected and aged for 8 days under the same rearing condition before they were flash-frozen in liquid nitrogen and stored at −80°C.

### Genome size estimation

We estimated genome size of the 5 karyotypes of interest using flow cytometry methods similar to those described in (Ellis, et al. 2014). Briefly, samples were prepared by using a 2mL Dounce to homogenize one head each from an internal control (*D. virilis* female, 1C=328 Mb) and one of the 5 karyotypes in Galbraith buffer (44mM magnesium chloride, 34mM sodium citrate, 0.1% (v/v) Triton X-100, 20mM MOPS, 1mg/mL RNAse I, pH 7.2). After homogenizing samples with 15-20 strokes, samples were filtered using a nylon mesh filter, and incubated on ice for 45 minutes in 25 ug/mL propidium iodide. Using a BD Biosciences LSR II flow cytometer, we measured 10,000 cells for each unknown and internal control sample. We ran samples at 10-60 events per second at 473 voltage using a PE laser at 488 nm. Fluorescence for each *D. melanogaster* karyotype was measured using the FACSDiva 6.2 software and recorded as the mode of the sample’s fluorescent peak interval. We calculated the genome size of the 5 karyotypes by multiplying the known genome size of *D. virilis* (328 Mb) by the ratio of the propidium iodide fluorescence in the unknown karyotype to the *D. virilis* control.

### Western blotting

We performed Western blots from acid-extracted histones, probing for H3K9me2, H3K9me3, H3K4me3, and total H3. Briefly, approximately 30 flies of each karyotype were dissected on dry ice to remove the abdomen. The resulting heads and thoraces were ground in PBS plus 10mM sodium butyrate, and were acid-extracted overnight at 4°C. Samples were then run on a 4-12% gradient bis-tris gel and transferred to a nitrocellulose membrane using Invitrogen’s iBlot Dry Transfer Device. After blocking with 5% milk in PBS, we incubated membranes overnight with either 1:1000 H3K9me2 antibody (Abcam ab1220), 1:2000 H3K9me3 antibody (Abcam ab8898), 1:2000 H3K4me3 antibody (Abcam ab8580), or 1:2000 H3 antibody (Abcam ab1791) in Hikari Signal Enhancer (Nacalai 02272). We then incubated membranes with 1:2500 secondary antibody (Licor 68070 and 32213), imaged bands on a Licor Odyssey CLx Imager, and quantified intensity using ImageJ.

### Chromatin-IP and sequencing

We performed ChIP-seq experiments using a standard protocol adapted from (Alekseyenko, et al. 2006). Briefly, approximately 2 mL of adult flash-frozen flies were dissected on dry ice, and heads and thoraces were used to fix and isolate chromatin. Following chromatin isolation, we spiked in 60uL of chromatin prepared from female *Drosophila miranda* larvae (approximately 1ug of chromatin); for replicate experiments, we used new preparations of *D. melanogaster* chromatin and the same *D. miranda* chromatin spike. We then performed immunoprecipitation using 4uL of one of the following antibodies: H3K9me2 (Abcam ab1220), H3K9me3 (Abcam ab8898), and H3K4me3 (Abcam ab8580).

After reversing the cross-links and isolating DNA, we constructed sequencing libraries using the BIOO NextFlex sequencing kit. Sequencing was performed at the Vincent J. Coates Genomic Sequencing Laboratory at UC Berkeley, supported by NIH S10 Instrumentation Grants S10RR029668 and S10RR027303. We performed 50bp single-read sequencing for our input and H3K4me3 libraries, and 100bp paired-end sequencing for the H3K9me2 and H3K9me3 libraries, due to their higher repeat content.

For H3K4me3, Pearson correlation values between the 5 karyotypes is very high, and the magnitude of difference between the samples is low (**Table S2**). For the two heterochromatin marks, Pearson correlation values between the two marks were generally high for all samples, and overlap of the top 40% of 5kb windows was similarly high for all samples (**Table S2**). Additionally, we obtained replicates for H3K9me3 for all samples except XX female, which has extremely high correlation values between H3K9me2 and H3K9me3. The unspiked replicate data for H3K9me3 correlate well with the *D. miranda* chromatin spike data that was used for the bulk of our analyses (**Table S2**).

We also generated replicate ChIP-seq data for H3K9me3 from XO, XXY, and XYY individuals using a different attached X stock, 4248, and a different ChIP-seq protocol, ULI-NChIP-seq, based on (Brind’Amour et al. 2015). Briefly, 4 flies from each of the three karyotypes were collected, heads were dissected, and along with a single *D. miranda* head, were homogenized in PBS and spun at 500g to isolate nuclei. MNase digestion was performed at 37°C for 5 minutes, at which point the reaction was stopped by the addition of 10% 100uM EDTA and incubated for 1 hr in complete immunoprecipitation buffer (20mM Tris-HCl pH 8.0, 2mM EDTA, 150mM NaCl, 0.1% Triton X-100, 1mM PMSF, 1x protease inhibitors). Samples were then incubated overnight at 4°C with 1ug of H3K9me3 antibody and 10ul of Dynabeads (Life Technologies 1006D). Libraries were then prepared using the BIOO NextFlex sequencing kit and sequenced at the Vincent J. Coates Genomic Sequencing Laboratory at UC Berkeley.

### RNA extraction and RNA-seq

We collected mated males and females of the various karyotypes, aged them for 8 days, and dissected and pooled 5 heads from each karyotype. A replicate set of individuals was collected from independent vials for the wildtype Canton-S, and independent crosses for the XO, XXY, and XYY individuals. We then extracted RNA and prepared stranded total RNA-seq libraries using Illumina’s TruSeq Stranded Total RNA Library Prep kit with Ribo-Zero ribosomal RNA reduction chemistry, which depletes the highly abundant ribosomal RNA transcripts (Illumina RS-122-2201). We performed 50bp single-read sequencing for all total RNA libraries at the Vincent J. Coates Genomic Sequencing Laboratory at UC Berkeley.

### Mapping of sequencing reads, and data normalization

For all *D. melanogaster* alignments, we used Release 6 of the genome assembly and annotation (Hoskins, et al. 2015). For all ChIP-seq datasets, we used Bowtie2 (Langmead and Salzberg 2012) to map reads to the genome, using the parameters “-D 15 –R 2 –N 0 –L 22 –i S,1,0.50 --no-1mm-upfront”, which allowed us to reduce cross-mapping to the *D. miranda* genome to approximately 2.5% of 50bp reads, and 1% of 100bp paired-end reads. We also mapped all ChIP-seq datasets to the *D. miranda* genome assembly (Ellison and Bachtrog 2013) to calculate the proportion of each library that originated from the spiked-in *D. miranda* chromatin versus the *D. melanogaster* sample. To correct for variable coverage based on GC content (GC content bias), we used a shell script written by (Flynn, et al. 2017) to calculate correction factors following (Benjamini and Speed 2012). In particular, we calculated the average coverage of uniquely mappable regions of the genome, binned by GC content in 5% intervals. These values correspond to the expected coverage across the genome based on GC content. To normalize coverage of repetitive elements based on GC content, we divided the observed coverage of the repeat by the expected coverage based on the GC content of the repetitive element. To normalize coverage of the genome by GC content, we divided the observed coverage by the expected coverage based on the GC content of the 5kb region.

To calculate ChIP signal, we first calculated the coverage across 5kb windows for both the ChIP and the input, and then normalized by the total library size, including reads that map to both *D. melanogaster* and the *D. miranda* spike. We then calculated the ratio of ChIP coverage to input coverage, and normalized by the ratio of *D. melanogaster* reads to *D. miranda* reads in the ChIP library, and then by the ratio of *D. melanogaster* reads to *D. miranda* reads in the input, to account for differences in the ratio of sample to spike present before immunoprecipitation. Note that this normalization procedure is accounts for differences in ploidy as well as genome size by using a ratio of ChIP coverage to input coverage (see **Figure S1**).

### Gene expression analysis

We first mapped RNA-seq reads to the ribosomal DNA scaffold in the Release 6 version of the genome, and removed all reads that mapped to this scaffold, as differences in rRNA expression are likely to be technical artifacts from the total RNA library preparation. We then mapped the remaining reads to the Release 6 version of the *D. melanogaster* genome using Tophat2 (Kim, et al. 2013), using default parameters. We then used Cufflinks and Cuffnorm to calculate normalized FPKMs for all samples. GO analysis was performed using GOrilla using ranked lists of differentially expressed genes (Eden, et al. 2009).

### Repeat libraries

We used two approaches to quantify expression of repeats. Our first approach was based on consensus sequences of known repetitive elements that were included in the Release 6 version of the *D. melanogaster* genome and are available on FlyBase. These included consensus sequences for 125 TEs. We also added the consensus sequences of three known satellite sequences, (Dodeca, Responder, and 359), to include larger non-TE repetitive sequences in our repeat analyses.

We were particularly interested in mis-regulation of the Y chromosome, which is poorly assembled. We therefore assembled repetitive elements *de novo* from male and female genomic DNA reads using RepARK (Koch, et al. 2014), setting a manual threshold for abundant kmers of 5 times the average genome coverage, which corresponds to a repetitive sequence occurring at least 5 times in the genome. To identify male-specific repeats, we mapped male and female genomic reads back to our *de novo* assembled repeats, and identified repeats that had high coverage in males and either no coverage or significantly lower coverage in females (**Figure S9**). After filtering in this way, we obtained 101 male-specific repeats comprising 13.7kb of sequence, with a median repeat size of 101bp.

