## Supplementary Material for "The Drosophila Y chromosome affects heterochromatin integrity genome-wide"

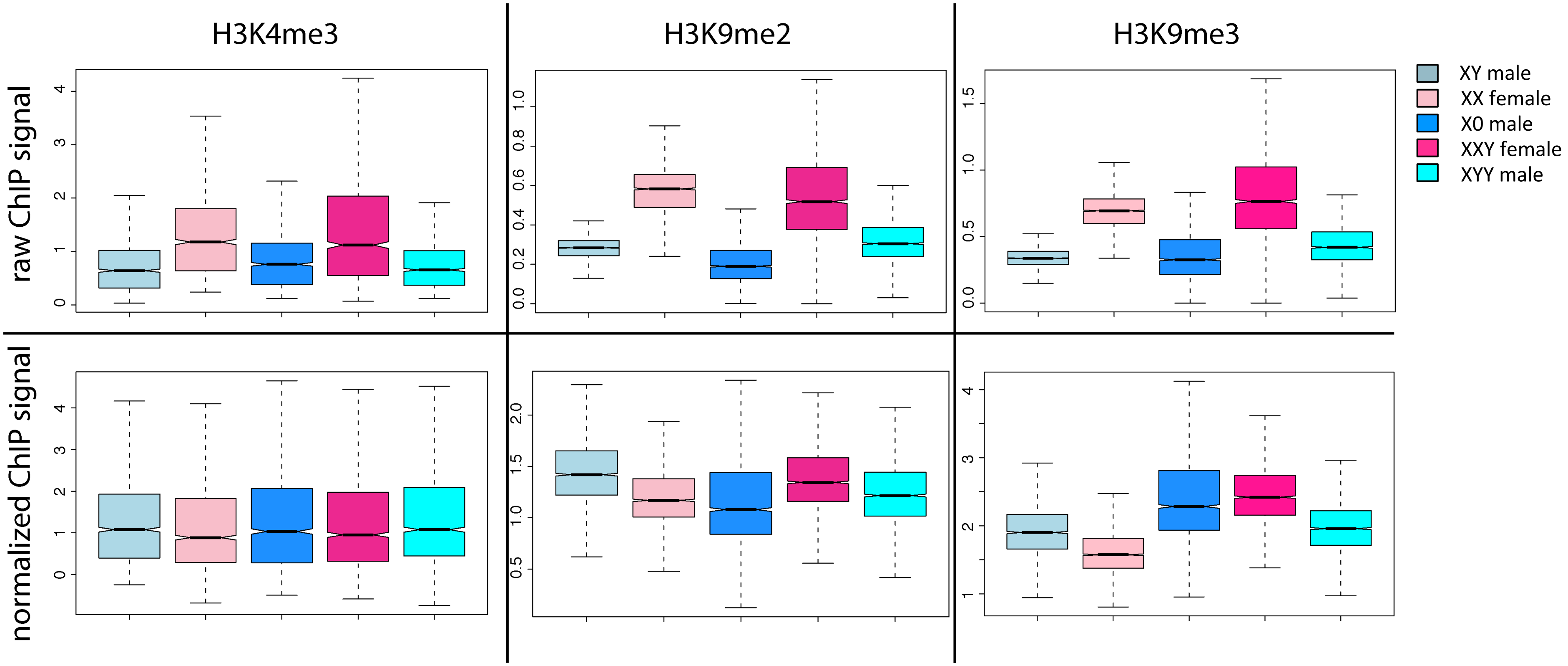


**Figure S1.** Normalization accounts for differences in ploidy of sex chromosomes. In the upper boxplots, we show the raw ChIP signal in genes on the X chromosome, where we know that there is a two-fold difference in ploidy between male and female karyotypes. In the lower boxplot, we plot the normalized signal in genes on the X chromosome, demonstrating that our normalization method corrects for differences in signal driven by differences in ploidy for all three histone modifications assayed.

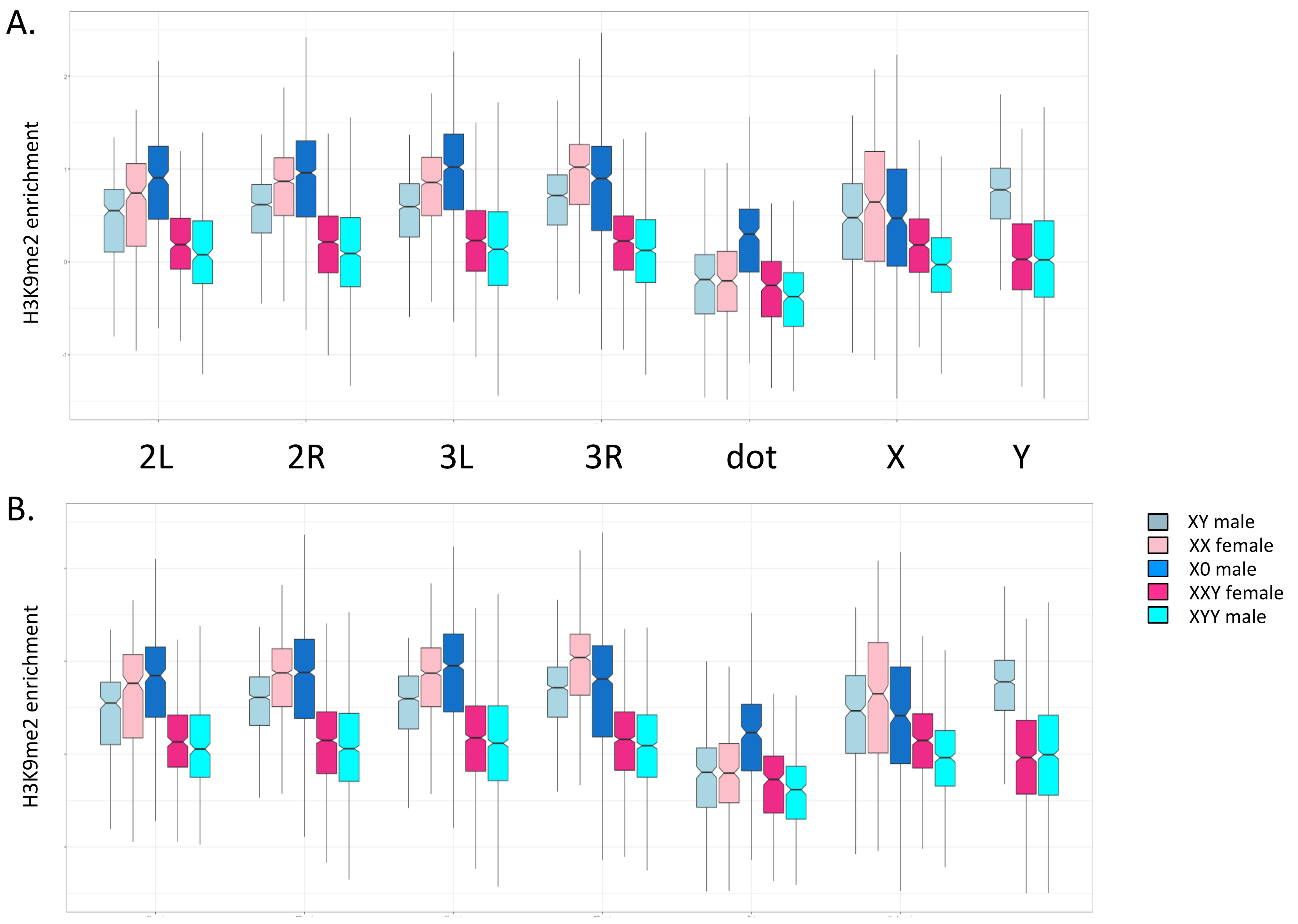


**Figure S2.** H3K9me2 enrichment using **A.** a normalization strategy based on a linear regression model **B.** only uniquely mapping reads at heterochromatic regions (pericentromere of chr2, chr3, X, the dot and Y chromosome). The box plots show the ChIP signal for all 5kb windows in different chromosomal regions, with boxes extending from the first to the third quartile and whiskers to the most extreme data point within 1.5 times the interquantile range.

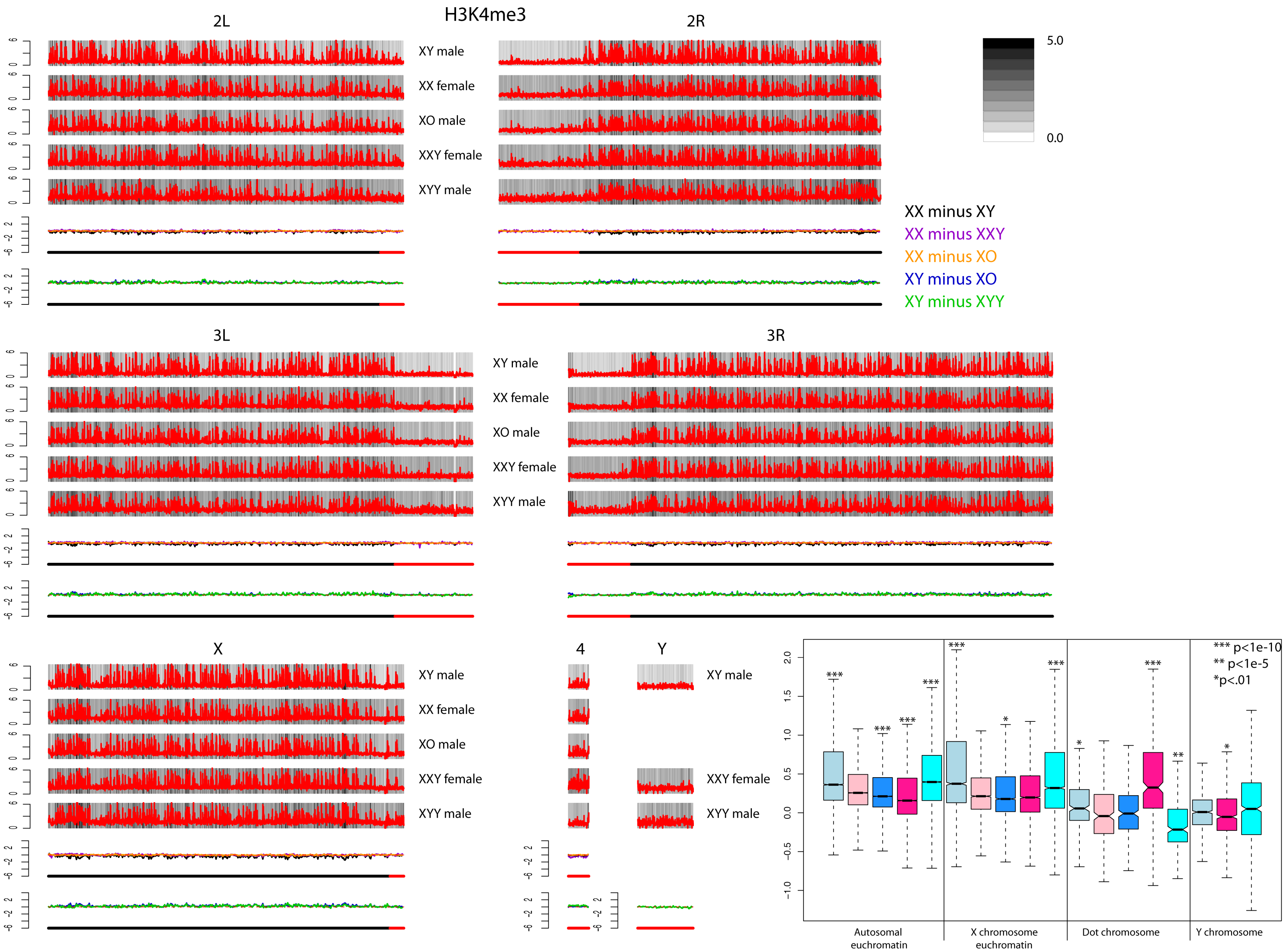


**Figure S3.** Genome-wide enrichment of H3K4me3 for *D. melanogaster* strains with different karyotypes along the different chromosome arms. These plots were made in the same manner as those for H3K4me3 (see **Figure 2**).

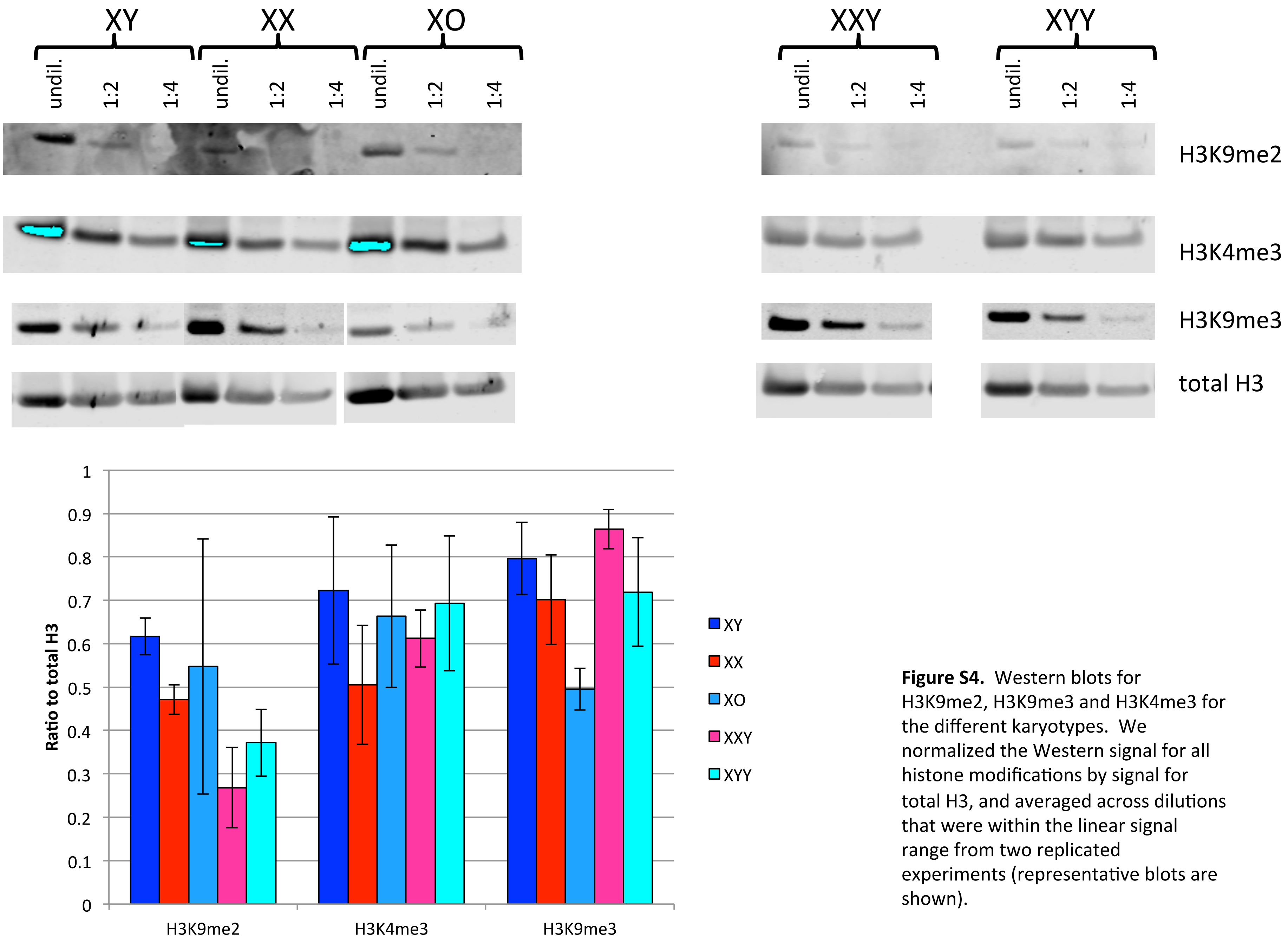


**Figure S4.** Western blots for H3K9me2, H3K9me3 and H3K4me3 for the different karyotypes. We normalized the Western signal for all histone modifications by signal for total H3, and averaged across dilutions that were within the linear signal range from two replicated experiments (representative blots are shown).

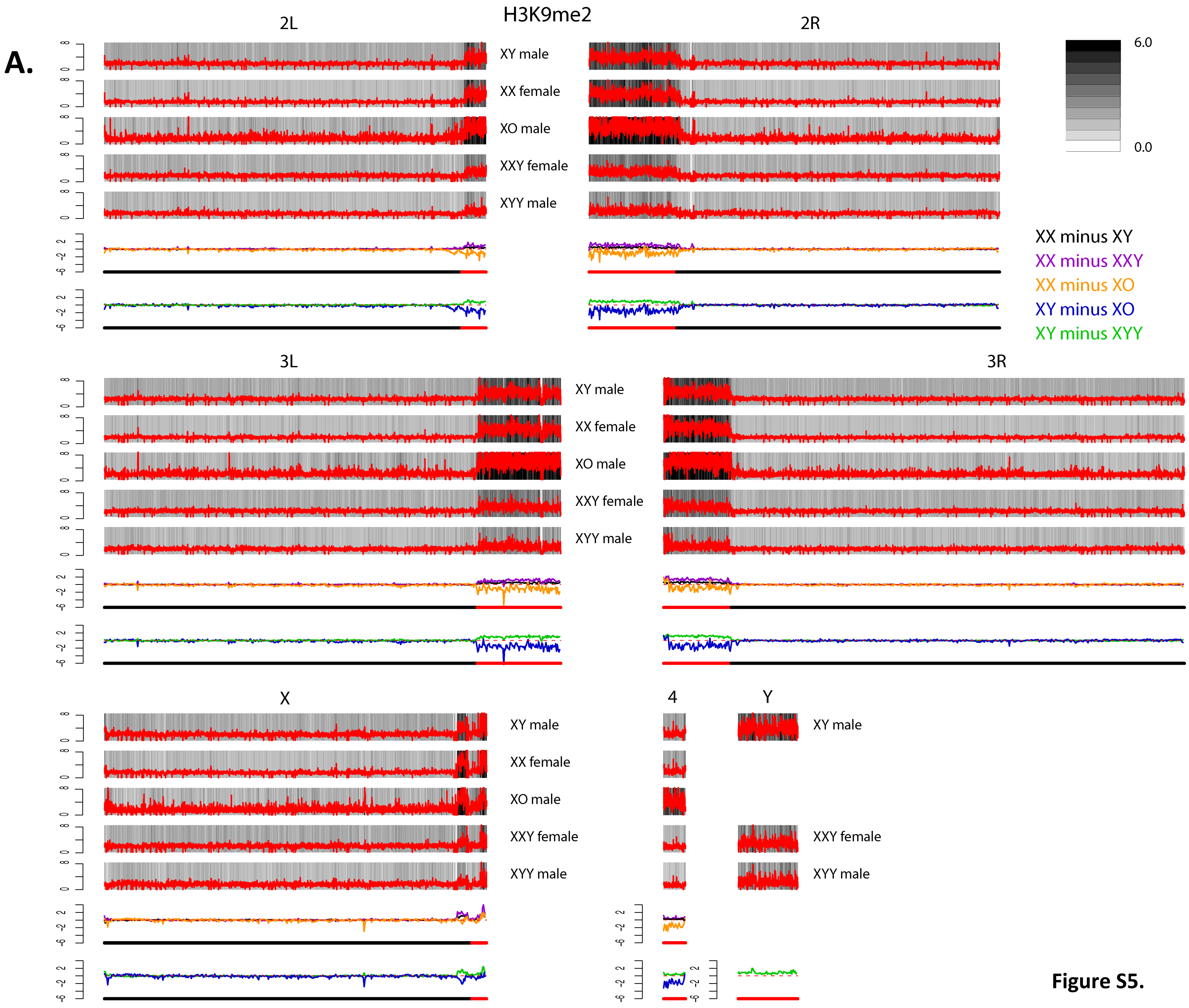
A.
H3K9me2
2L
2R
XY male
XX female
XO male
XXY female
XYY male
6.0
0.0
XX minus XY
XX minus XXY
XX minus XO
XY minus XO
XY minus XYY
3L
3R
XY male
XX female
XO male
XXY female
XYY male
X
4
Y
XY male
XX female
XO male
XXY female
XYY male
XY male
XXY female
XYY male

**Figure S5.**

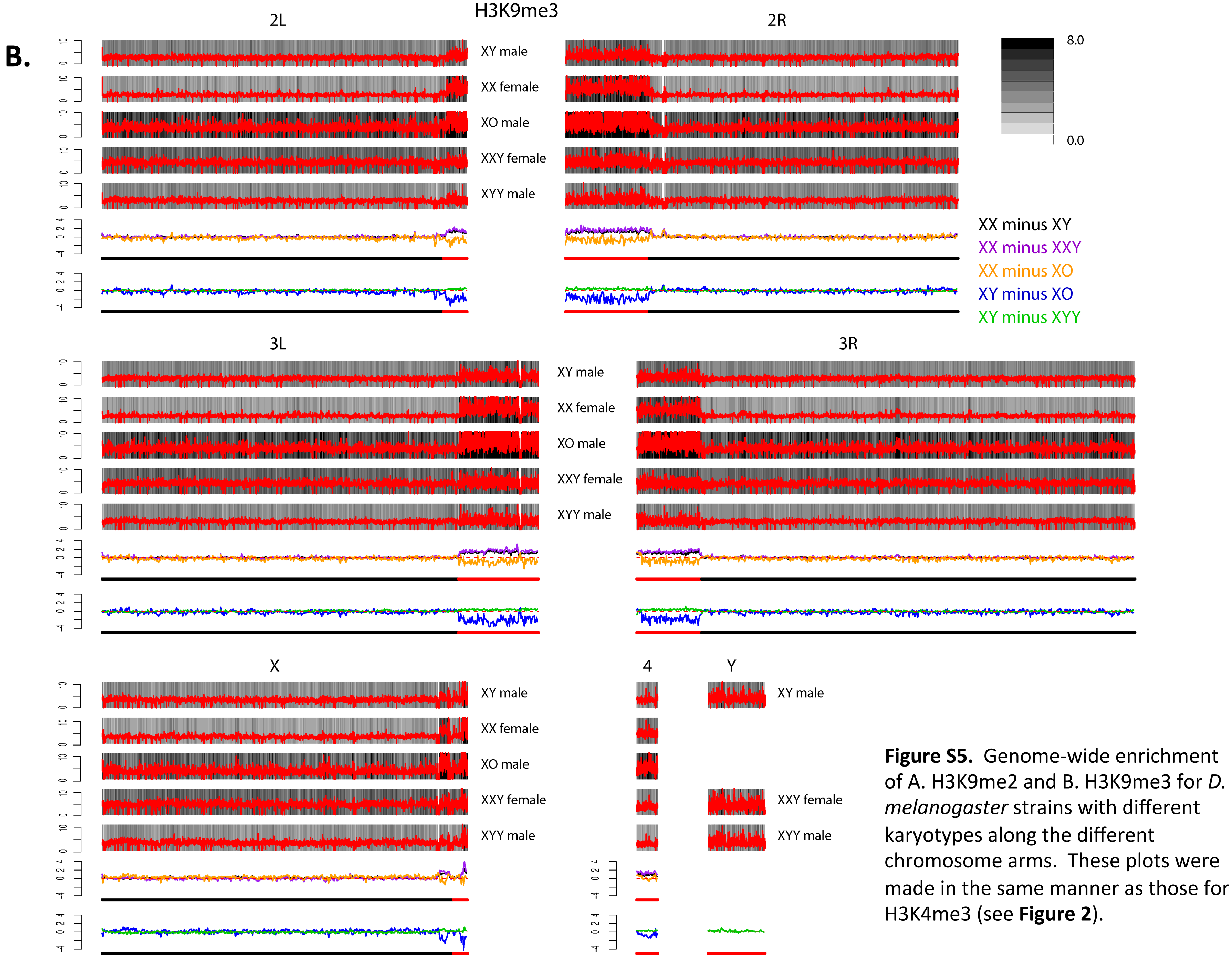


**Figure S5.** Genome-wide enrichment of A. H3K9me2 and B. H3K9me3 for *D. melanogaster* strains with different karyotypes along the different chromosome arms. These plots were made in the same manner as those for H3K4me3 (see **Figure 2**).

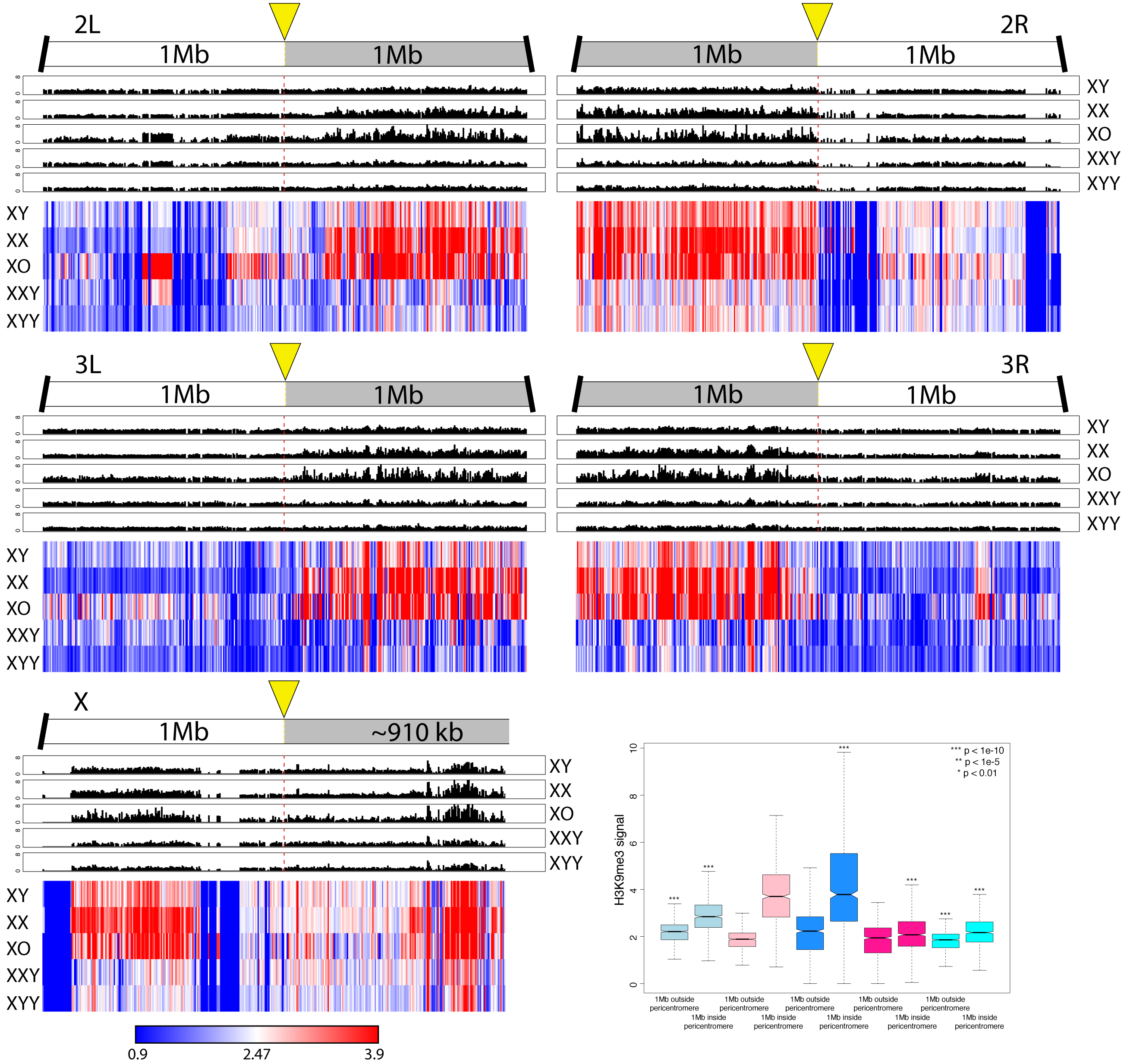


**Figure S6.** Enrichment of H3K9me3 within 1Mb of the heterochromatin/ euchromatin boundaries. The plots were made in the same manner as for H3K9me2 (see **Figure 5**).

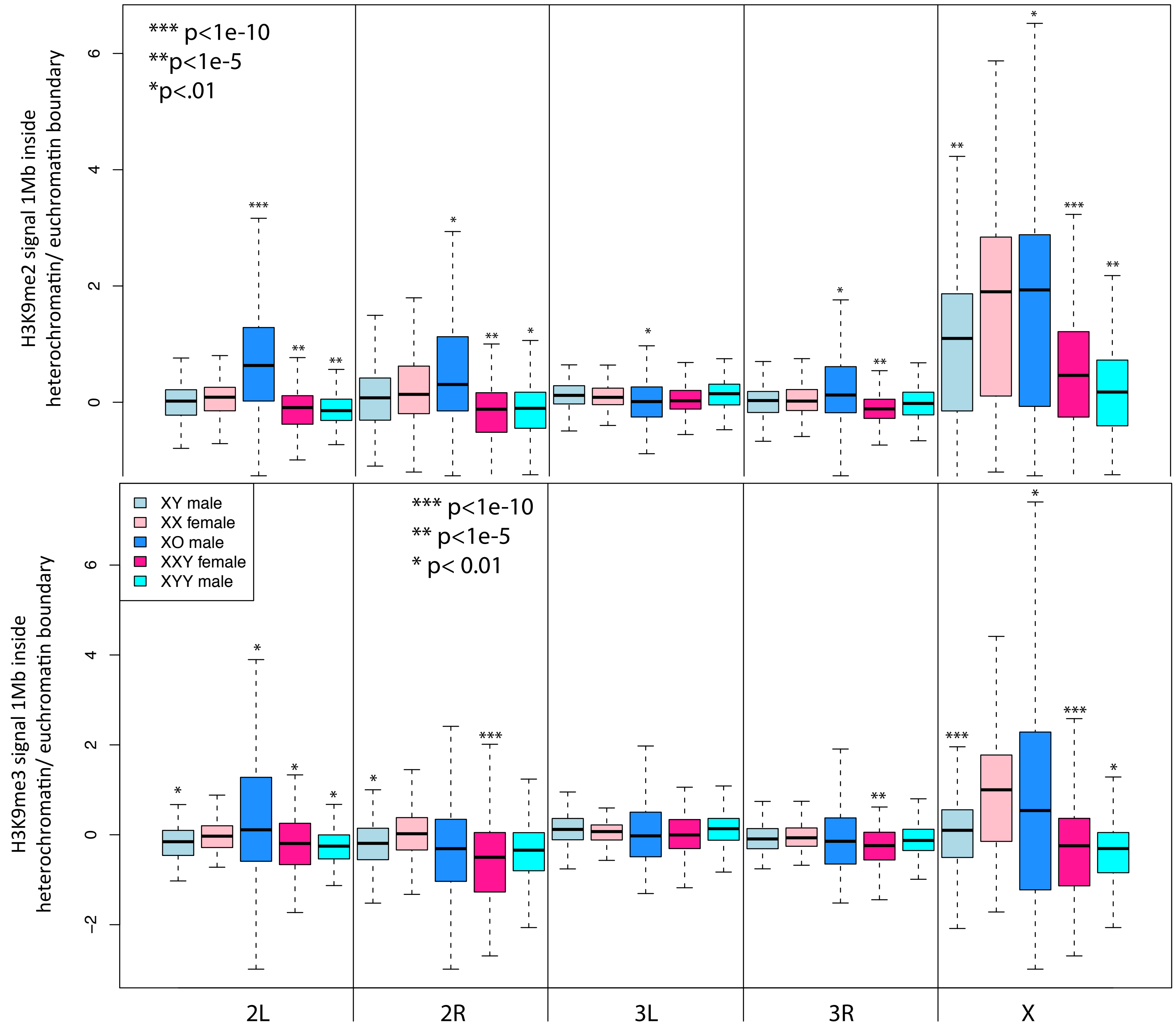


**Figure S7.** Normalized H3K9me2 and H3K9me3 signal of 5kb windows in euchromatic regions within 1Mb of the heterochromatin/ euchromatin boundary by chromosome arm. Boxes extend from the first to the third quartile and whiskers to the most extreme data point within 1.5 times the interquantile range. Significance values for XY and XXY individuals were calculated relative to XX females, and significance values for XO and XYY individuals were calculated relative to XY males, using the Wilcoxon test.

A.

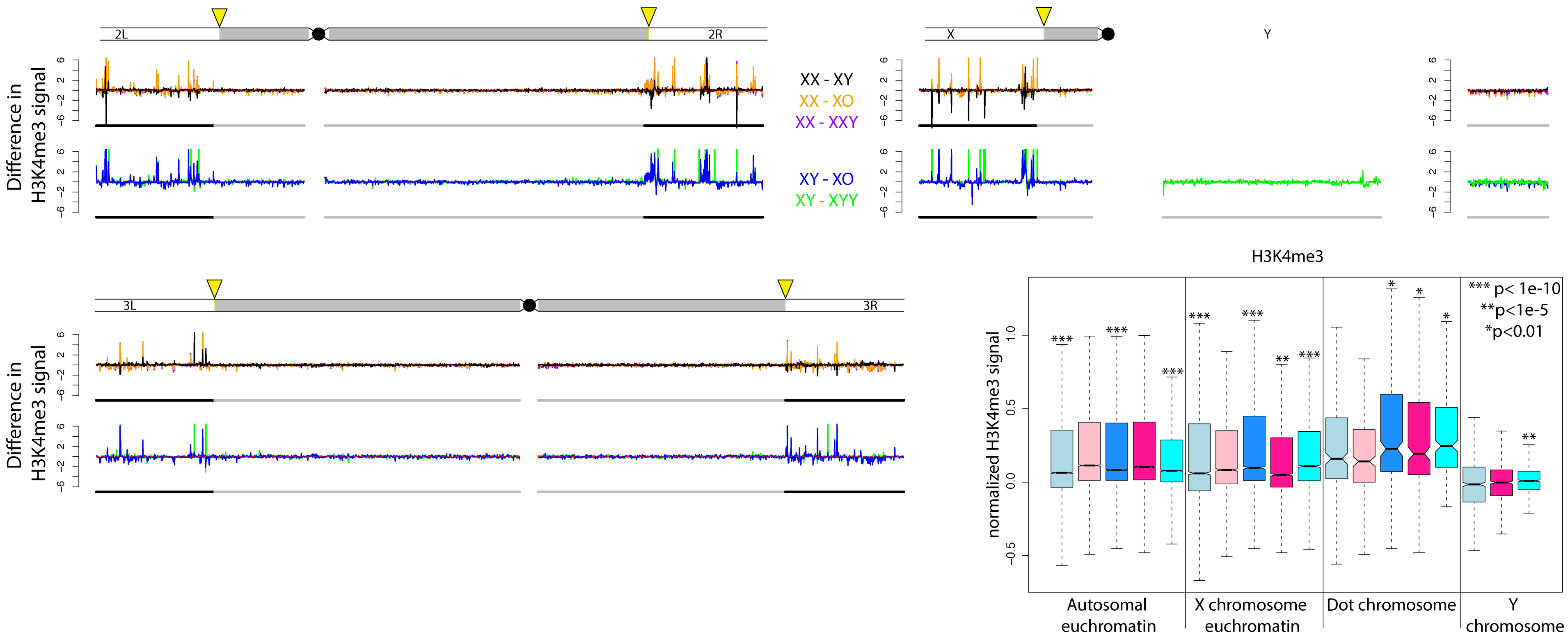


**Figure S8.** Biological replicate for **A.** H3K4me3 and B. H3K9me3 (sample without a spike).

B.

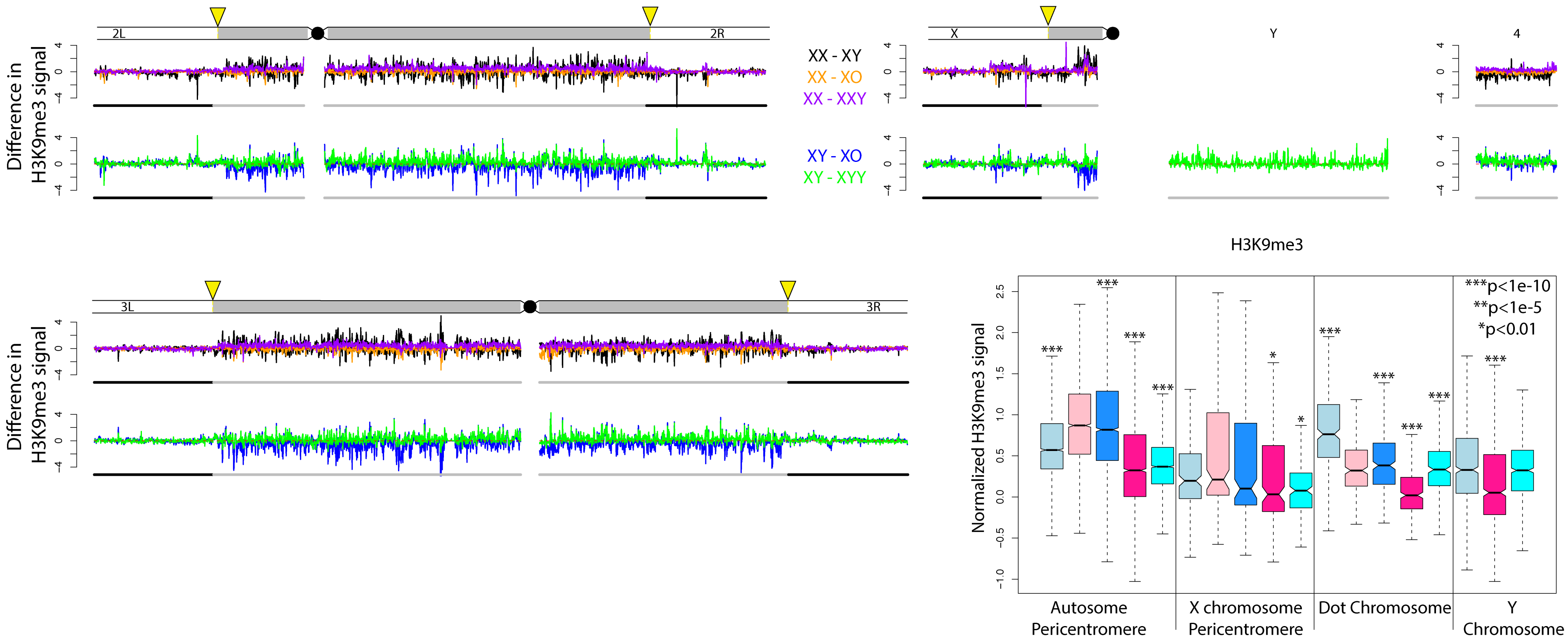


**Figure S8.** Biological replicate for **A.** H3K4me3 and B. H3K9me3 (sample without a spike).

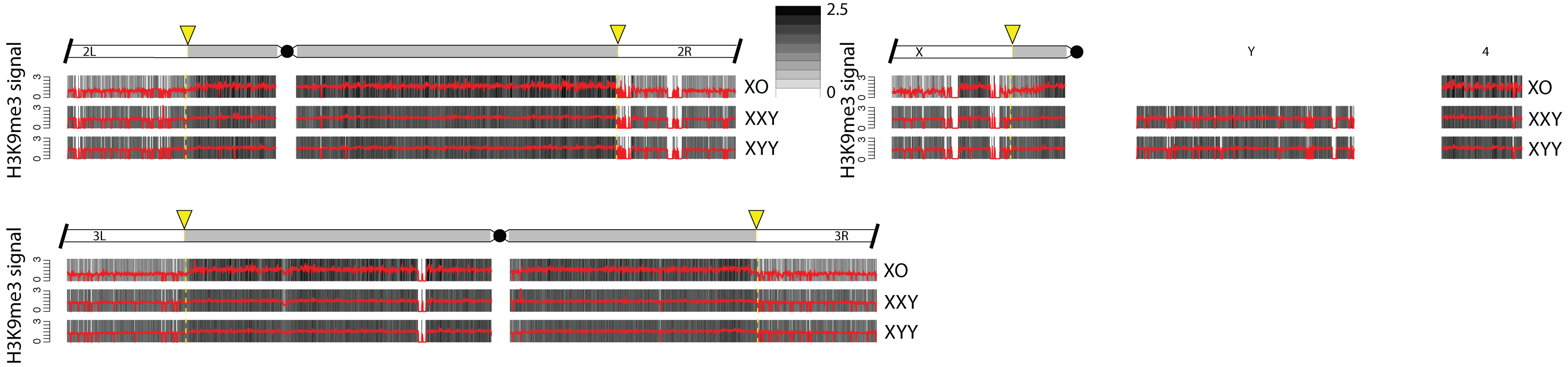


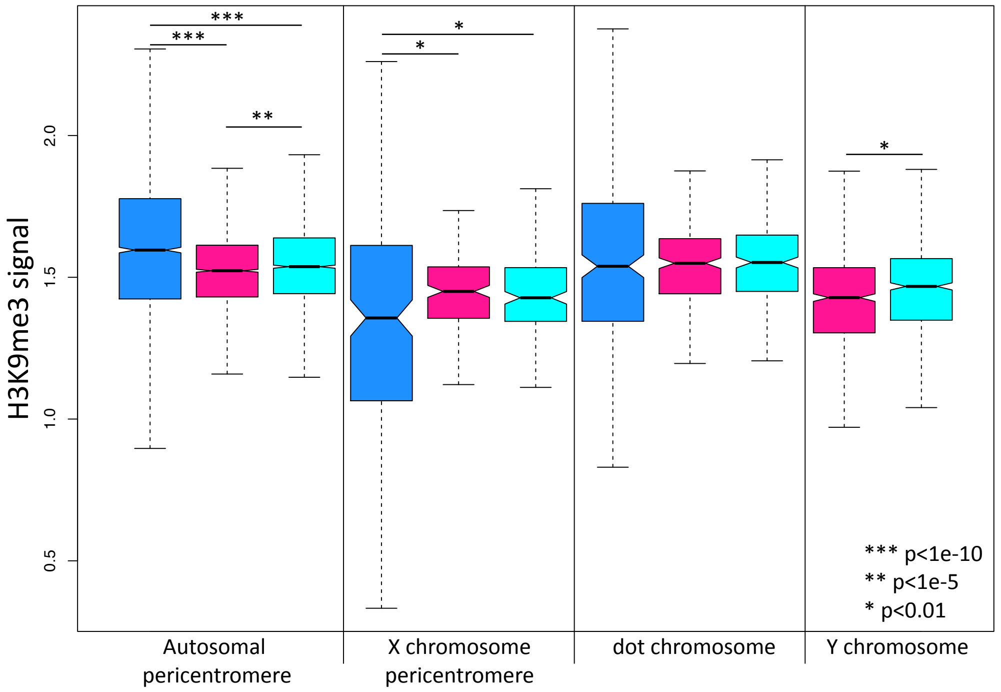


**Figure S9.** H3K9me3 ChIPs for replicate karyotypes with aberrant sex chromosomes. X0/XXY and XYY flies were generated by crossing Canton-S flies with the 4248 stock (a stock with a different attached-X, attached X-Y chromosome). For legend see Fig. 2-4.

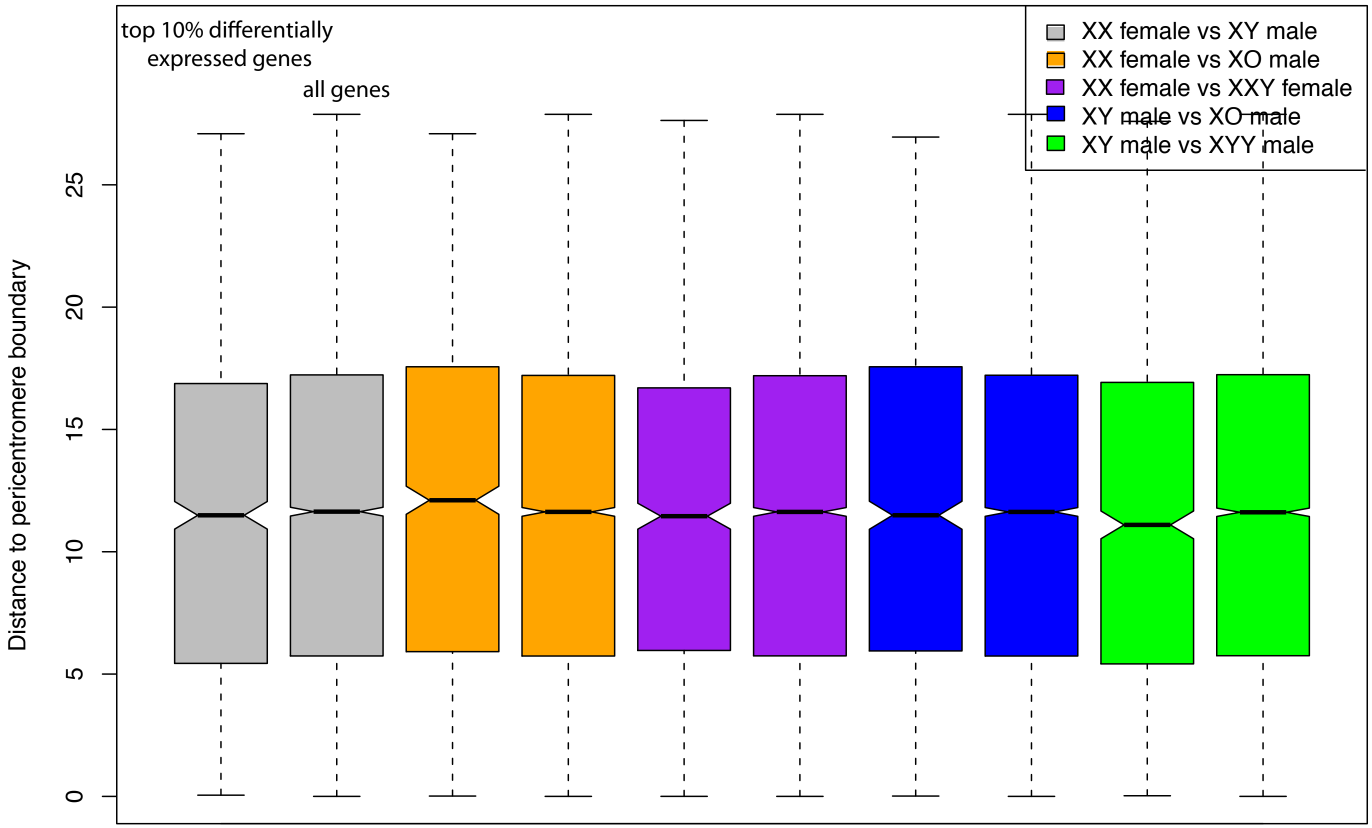


**Figure S10.** Distance to the pericentromere boundary (in Mb) of the top 10% of differentially expressed genes compared to all genes. For each pair of boxplots, the top 10% of differentially expressed genes are on the left, and all genes are on the right. Boxes extend from the first to the third quartile and whiskers to the most extreme data point within 1.5 times the interquantile range. None of the comparisons are significant (Wilcoxon test).

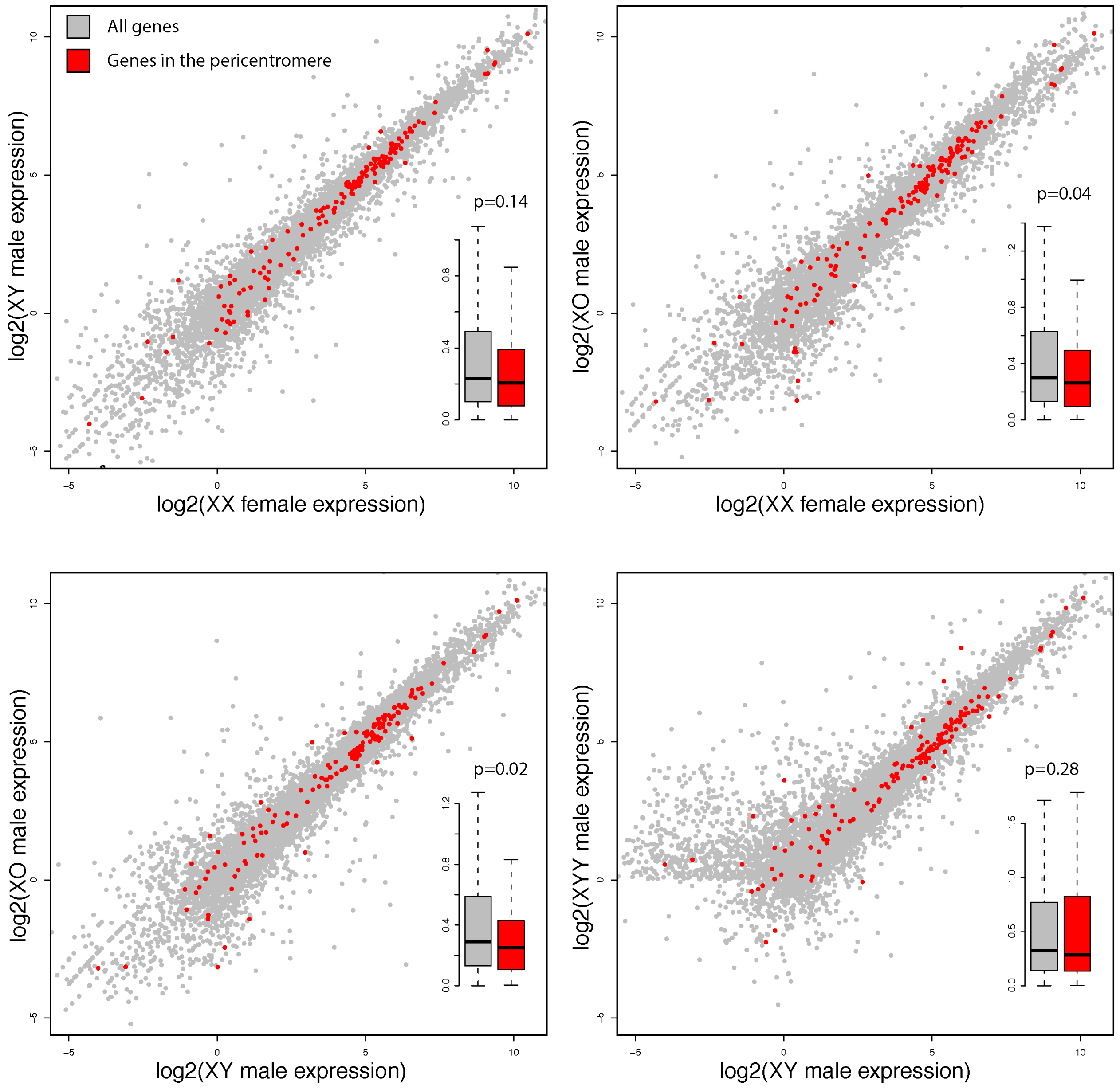


**Figure S11**. Pairwise comparisons of gene expression for different karyotypes, by location. Genes marked in red are located in the pericentromere, and all other genes are in gray. Boxplots summarize the absolute fold change (on a log2 scale) in expression between the two karyotypes, and p-values were calculated using the Wilcoxon test.

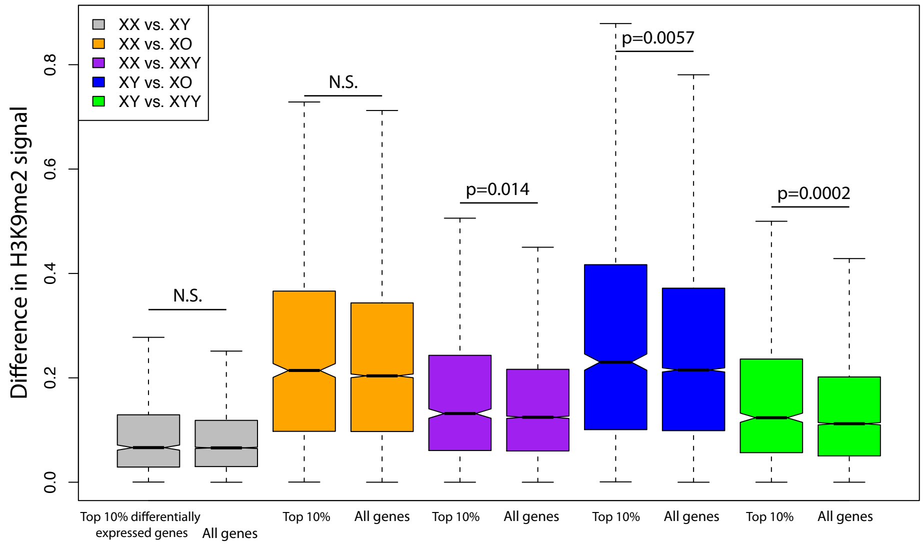


difference in H3K9me2 signal of genes differentially expressed vs. all genes

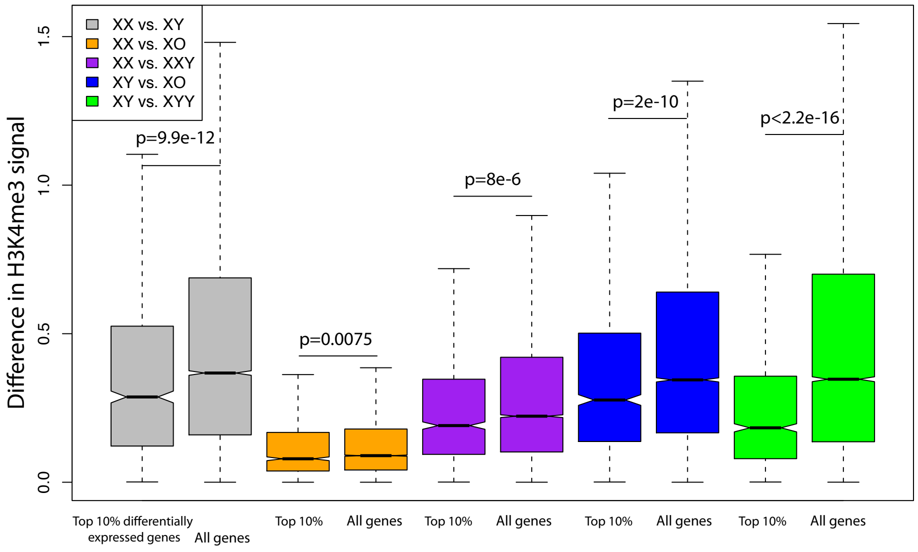


difference in H3K4me3 signal of genes differentially expressed vs. all genes

**Figure S12.** Difference in signal of H3K9me2 and H3K4me3 in the top 10% of differentially expressed genes ("Top 10%") compared to all genes for different pairwise comparisons of karyotypes. P-values were calculated for all comparisons using the Wilcoxon test.

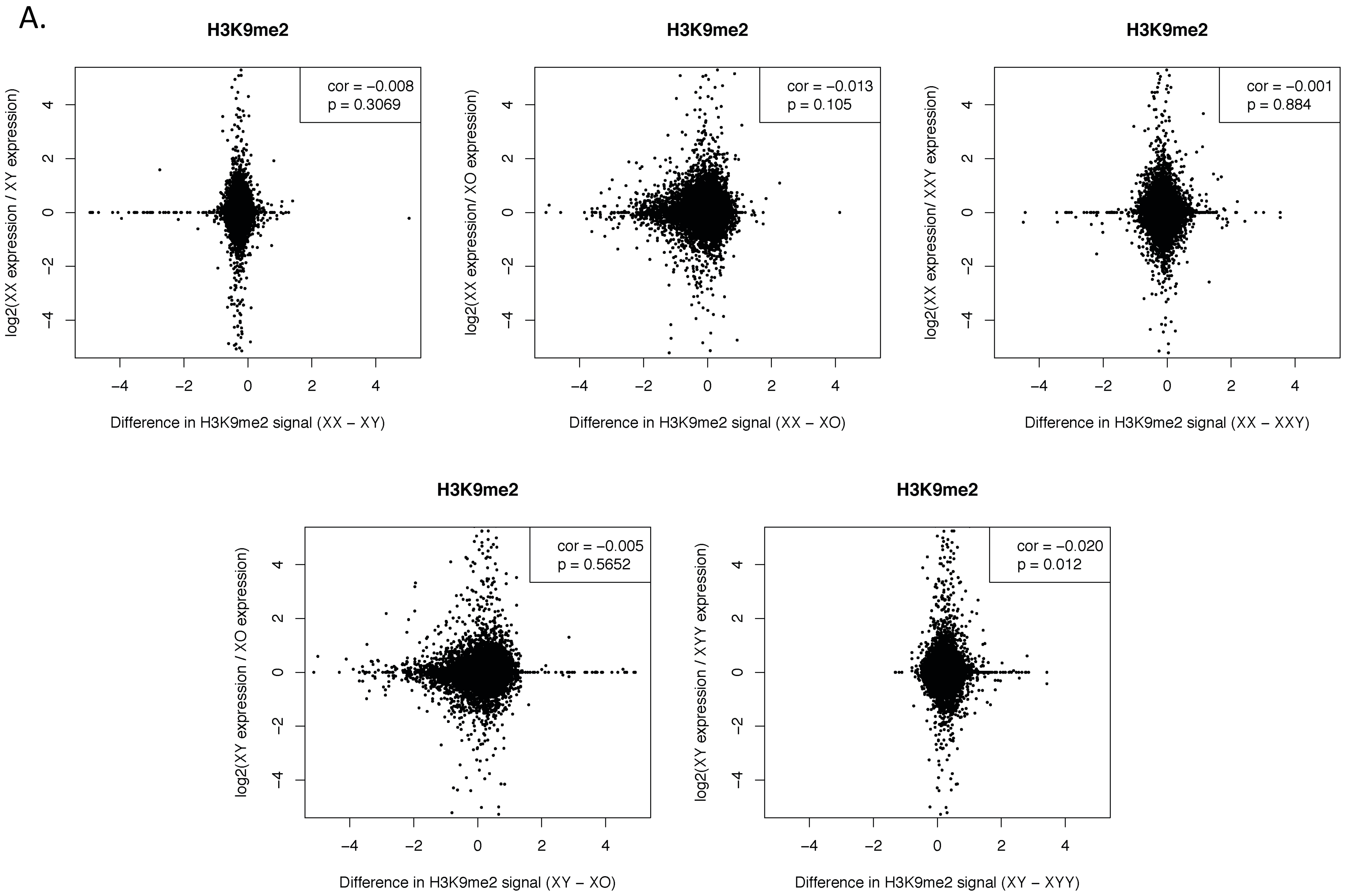


**Figure S13.** Correlations between expression differences and heterochromatin differences based on **A.** H3K9me2 or **B.** H3K9me3 ChIPs for different karyotypes.

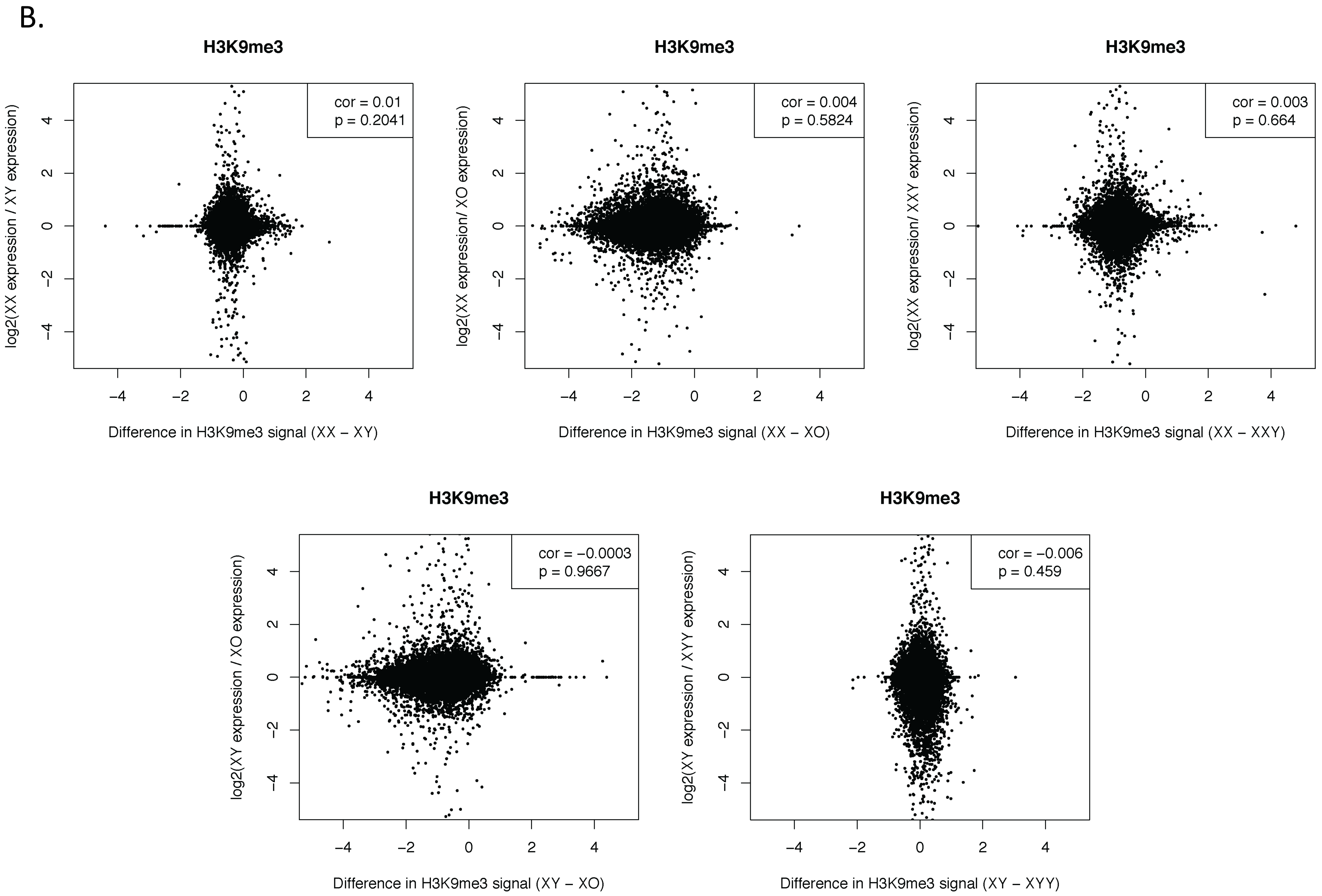


**Figure S13.** Correlations between expression differences and heterochromatin differences based on **A.** H3K9me2 or **B.** H3K9me3 ChIPs for different karyotypes.

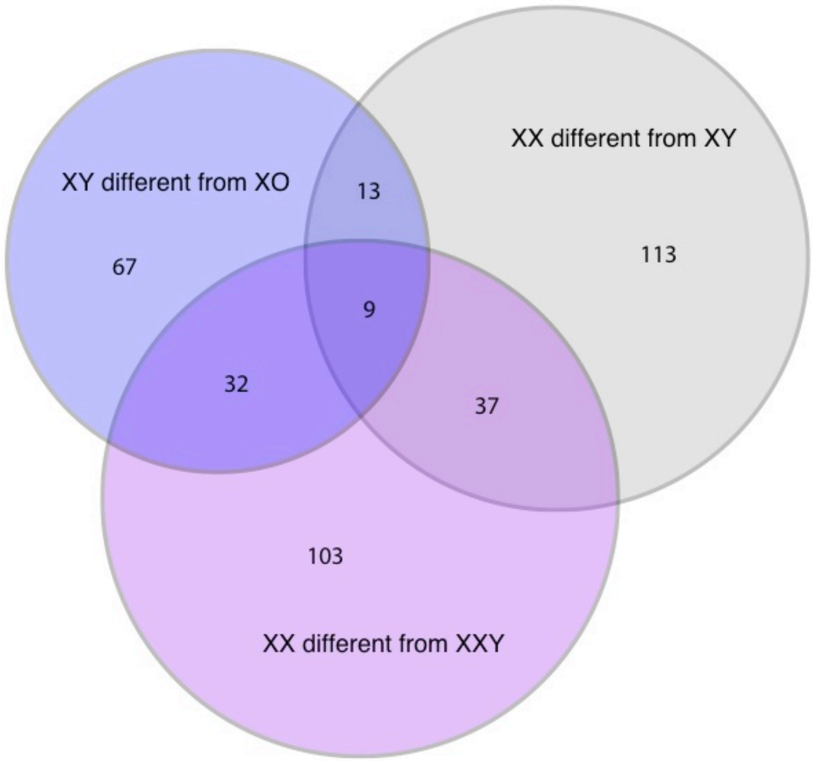


**Figure S14.** Overlap of genes categorized as significantly differently expressed by Cuffdiff between wildtype XY male and XX female, XX vs. XXY females, and XY vs. XO males.

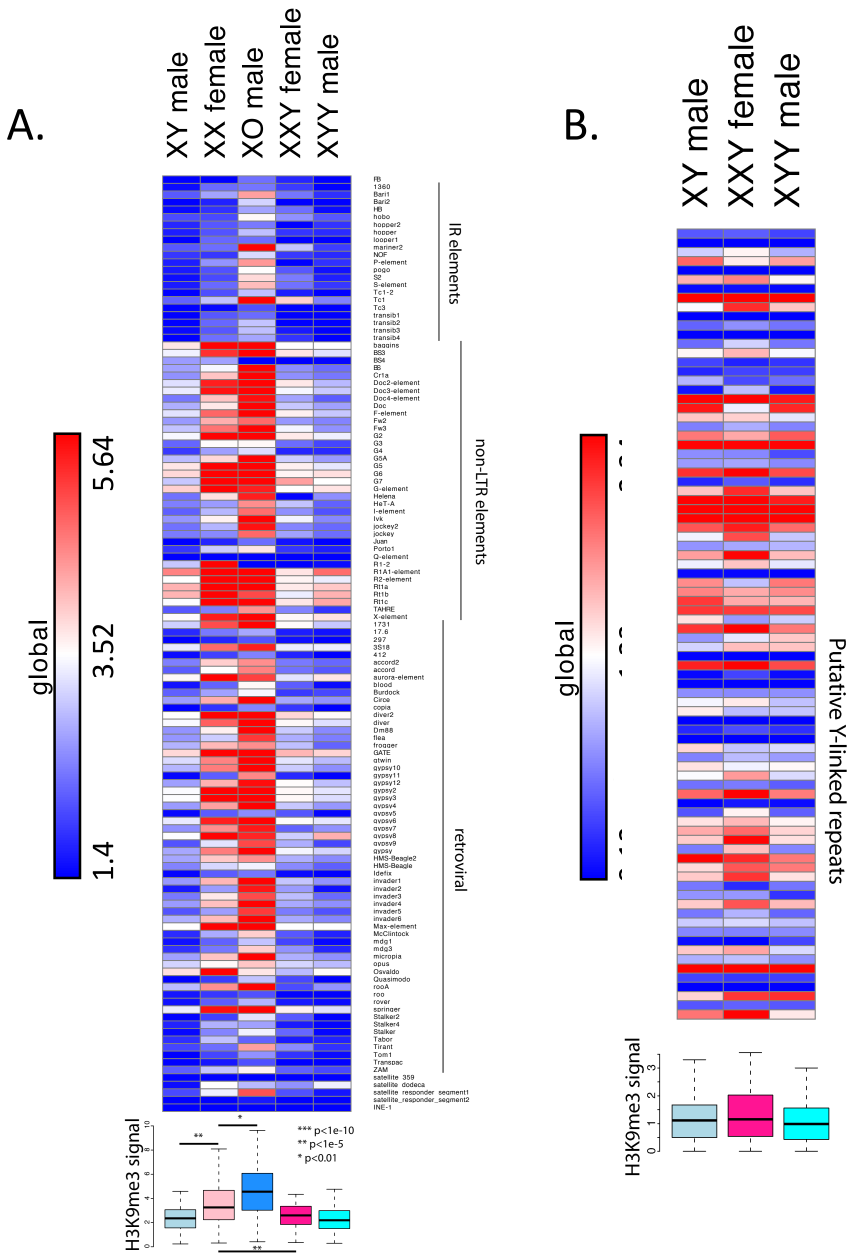


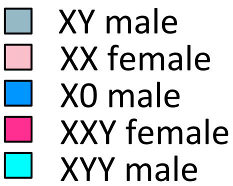


**Figure S15.** Enrichment of H3K9me3 at different TE families. **A.** All repeats from the library of consensus transposable elements and satellites from FlyBase. Boxplots summarize H3K9me3 signal in these consensus repeats, and p-values were calculated using the Wilcoxon test. **B.** Putatively Y-linked (male-specific) *de novo* assembled repeats only. Boxplots summarize H3K9me3 signal in putatively Y-linked repeats, and p-values were calculated using the Wilcoxon test.

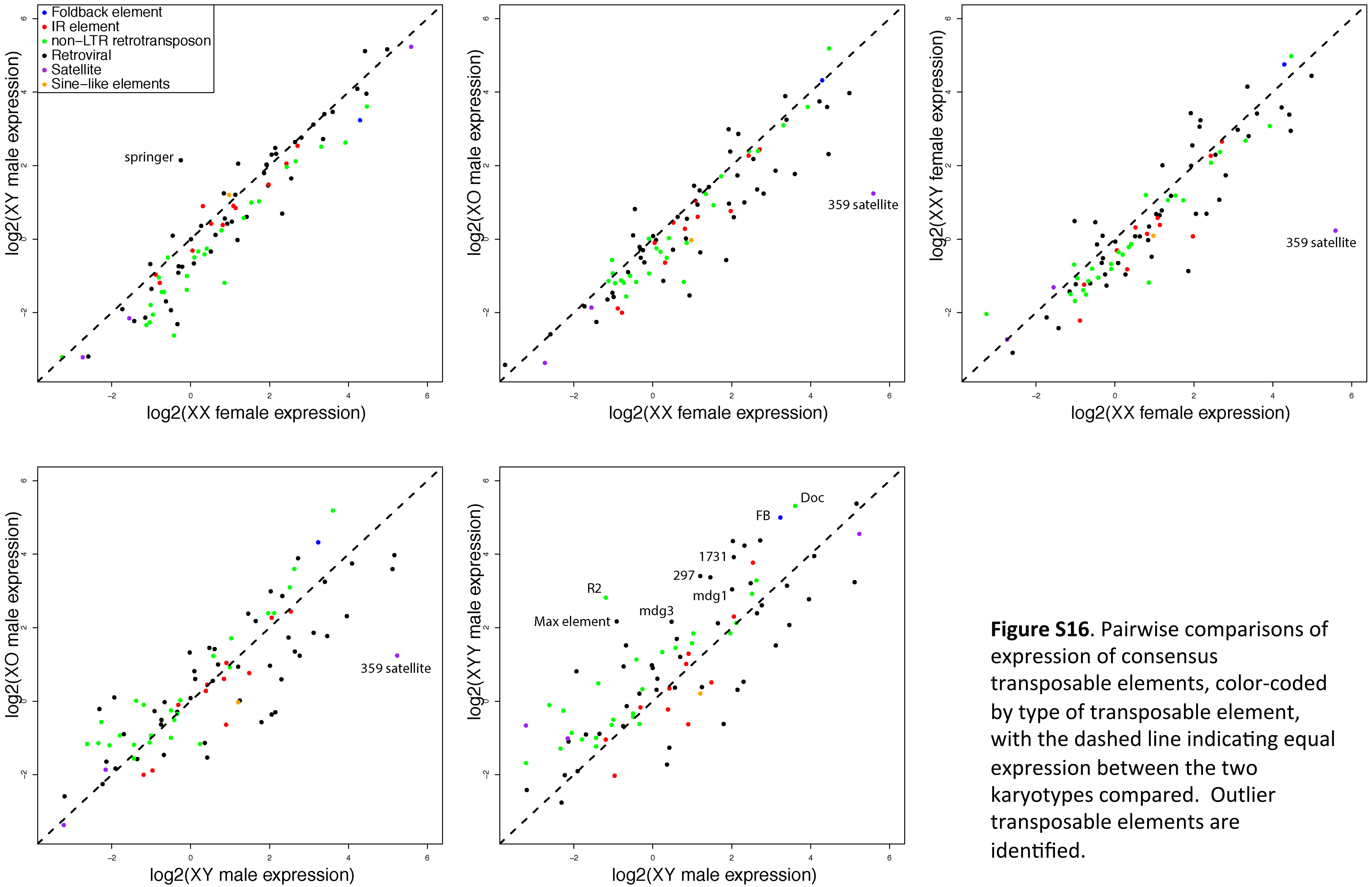


**Figure S16**. Pairwise comparisons of expression of consensus transposable elements, color-coded by type of transposable element, with the dashed line indicating equal expression between the two karyotypes compared. Outlier transposable elements are identified.

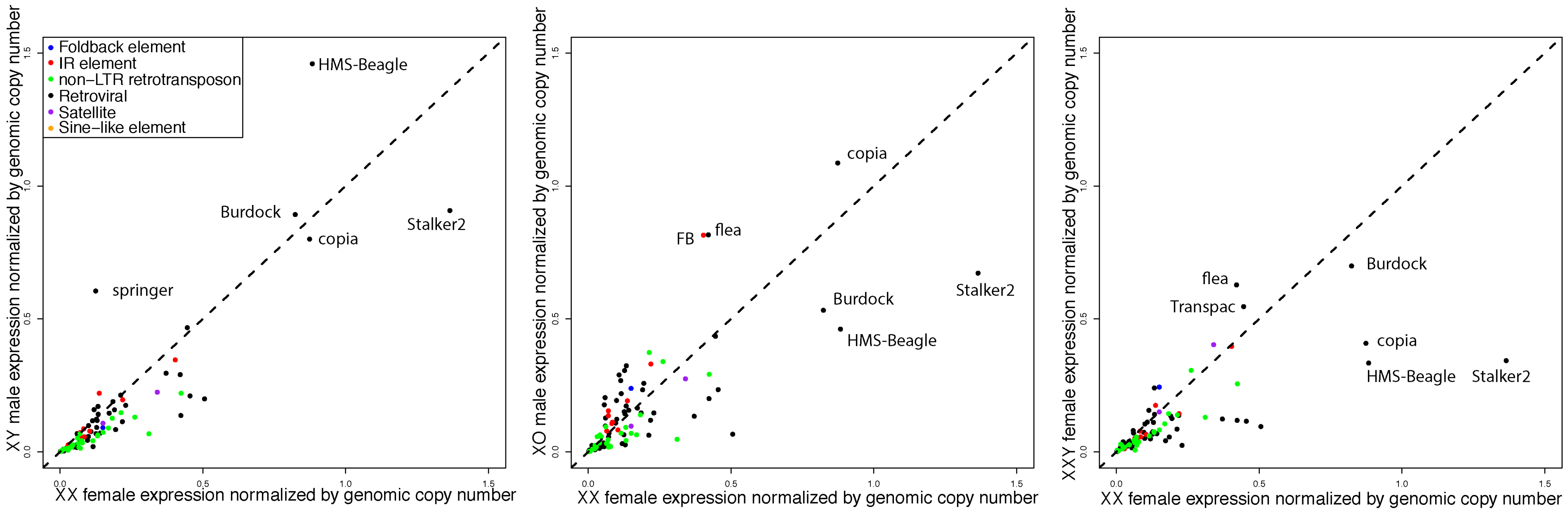


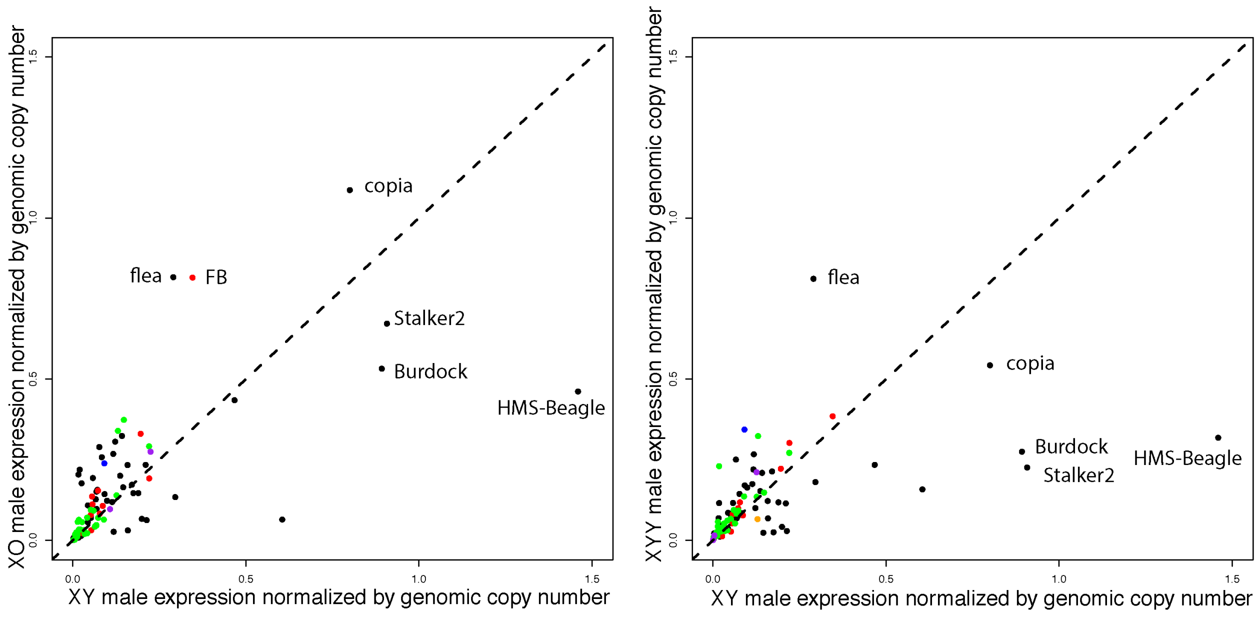


**Figure S17**. Pairwise comparisons of expression of consensus transposable elements normalized by genomic copy number, as estimated by coverage of each element from genomic DNA sequences, color-coded by type of transposable element. The dashed line indicates equal normalized expression between the two karyotypes compared. Outlier transposable elements are identified.

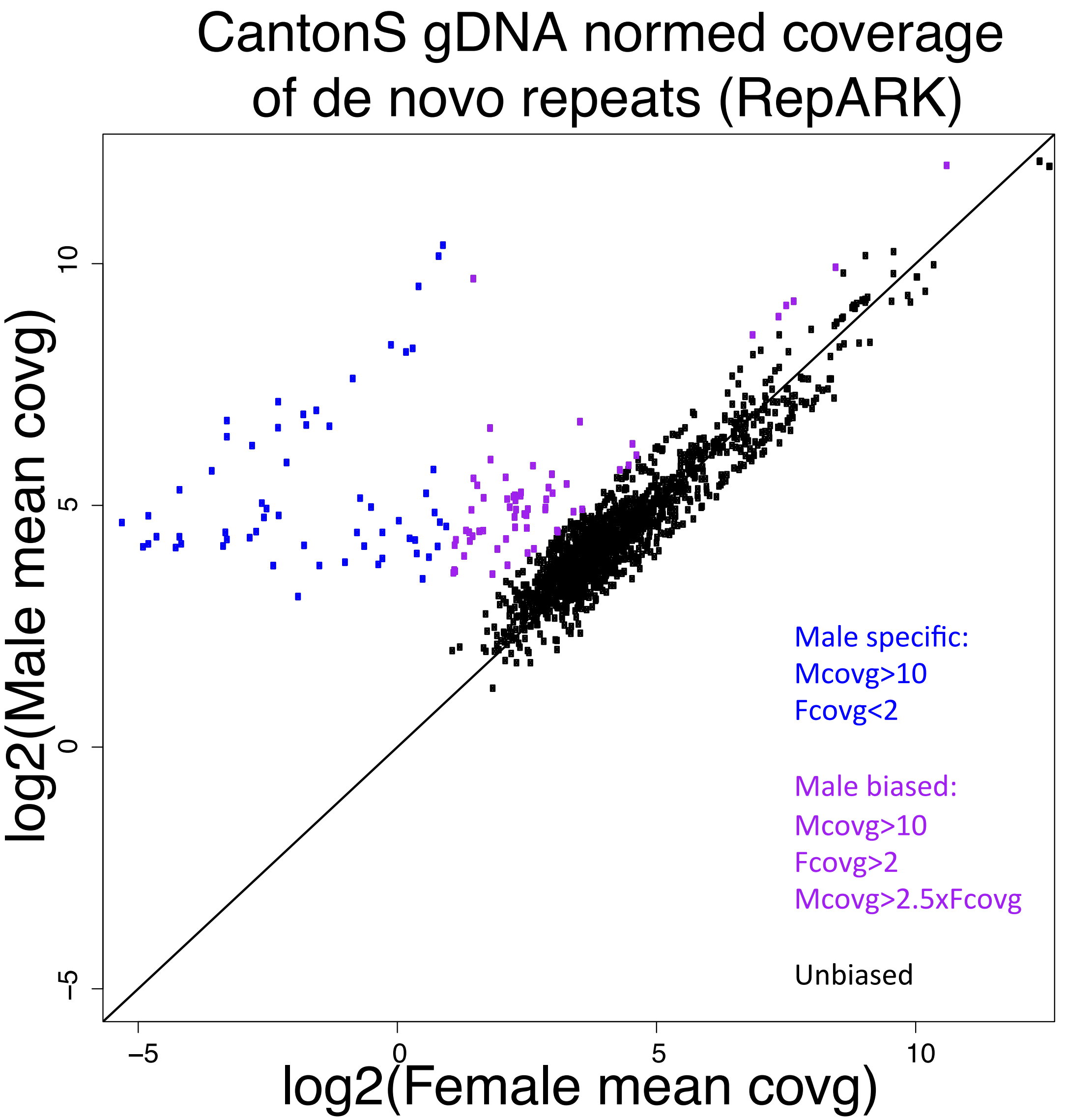


**Figure S18.** Categorization of *de novo* assembled repeats from RepARK as male-specific, male-biased, or unbiased based on coverage of female and male genomic reads. After *de novo* assembling repeats, we mapped male and female genomic reads to the repeats and removed all repeats that did not have at least 5 times the average genome coverage in one sex. We categorized male-specific repeats as those with at least 10 times the average male genome coverage and less than 2 times the average female genome coverage, and male-biased repeats as those with at least 10 times the average male genome coverage, at least 2 times the average female genome coverage, and male coverage at least 2.5 times higher than female coverage.

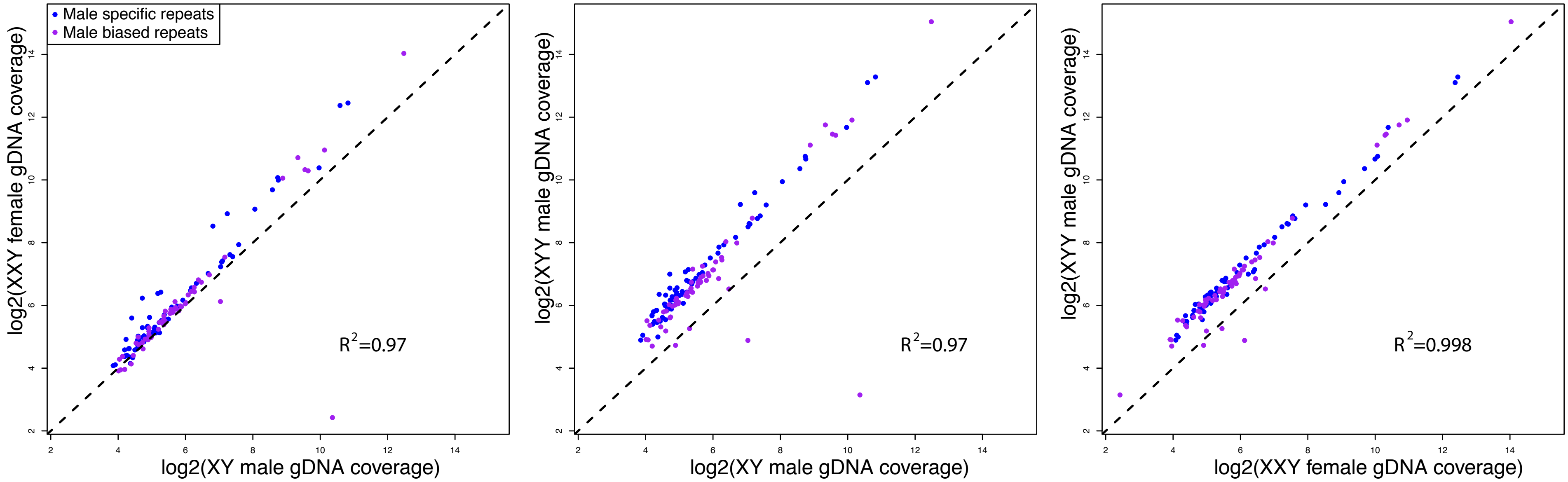


**Figure S19.** Pairwise comparisons of genomic coverage of *de novo* assembled male-biased and male-specific repeats for karyotypes containing at least one Y chromosome (XY, XXY, and XYY). Genomic coverage is estimated from sequencing reads of libraries constructed from genomic DNA of XY, XXY and XYY flies. The dashed line indicates equal coverage of repeats in the karyotypes compared.

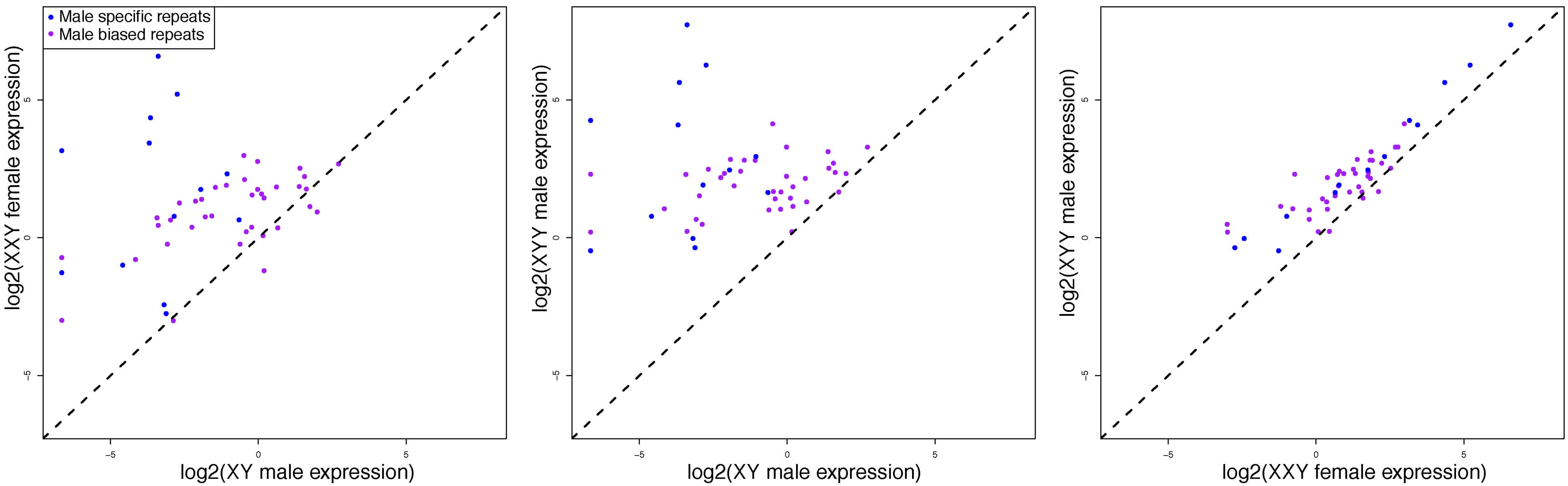


**Figure S20**. Pairwise comparisons of expression of *de novo* assembled male-biased and male-specific repeats for karyotypes containing at least one Y chromosome (XY, XXY, and XYY). Expression levels are not corrected by genomic copy number, instead reflecting the total number of transcripts from putatively Y-linked repeats in each karyotype.

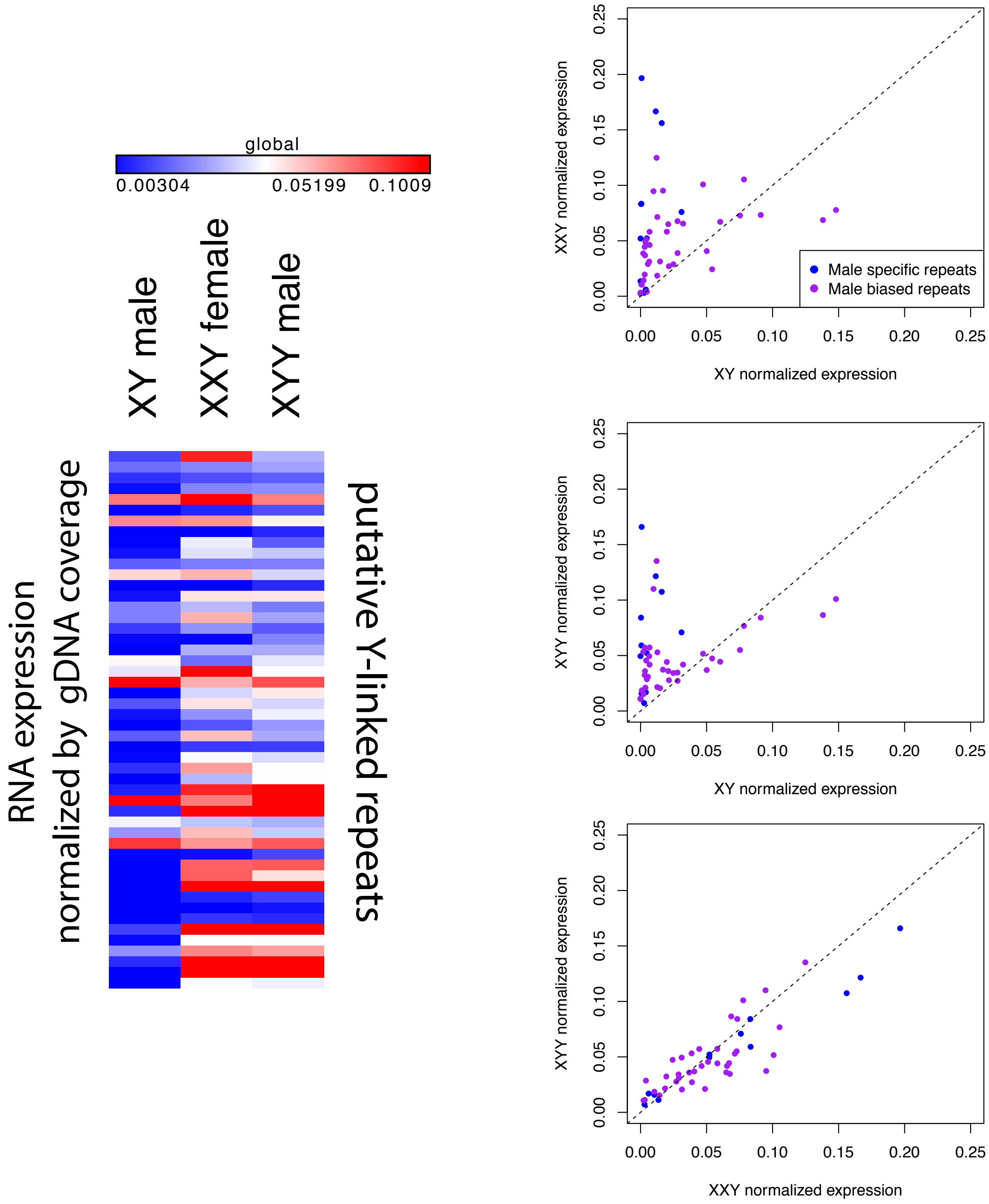


**Figure S21**. Expression of male-specific/ male-biased repeats normalized by genomic copy number in karyotypes with at least one Y chromosome (XY, XXY, and XYY). The heatmap indicates the $90^{th}$ percentile or higher values in deep red, the $10^{th}$ percentile or lower values in deep blue, across all karyotypes and repeats. The scatterplots show pairwise comparisons of expression values normalized by genomic coverage, with the dashed line indicating equal expression between the two karyotypes compared.

**Table S1**. Flow cytometry estimates of genome sizes for flies with different karyotypes. Shown are the measurements from 3 independent replicates.

| | Replicate 1 | Replicate 2 | Replicate 3 | Mean | SE |
|---|---|---|---|---|---|
| **CantonS F** | 177.6 | 177.2 | 180.4 | 178.4 | 1.8 |
| **CantonS M** | 179.9 | 178.0 | 178.5 | 178.8 | 1.0 |
| **XO** | 159.7 | 157.9 | 161.3 | 159.6 | 1.7 |
| **XXY** | 195.8 | 192.8 | 195.7 | 194.8 | 1.7 |
| **XYY** | 194.5 | 200.5 | 197.4 | 197.4 | 3.0 |

**Table S2**. Pearson correlation coefficients of signal of *D. miranda* spike for the H3K4me3, H3K9me2 and K3K9me3 ChIP's. The signal is calculated as the ratio of reads from the immunoprecipitation (normalized by library size) to reads from the input (normalized by library size) and multiplied by the normalization factor, as described in the methods. Correlation coefficients are calculated based on normalized ChIP-seq signal in 5-kb non-overlapping windows. Values below the diagonal are without GC correction, and values above the diagonal are with the GC correction.

**H3K4me3**

| | XY | XX | X0 | XXY | XYY |
|---|---|---|---|---|---|
| XY | | 0.87 | 0.97 | 0.91 | 0.92 |
| XX | 0.87 | | 0.89 | 0.9 | 0.9 |
| X0 | 0.82 | 0.96 | | 0.91 | 0.94 |
| XXY | 0.82 | 0.71 | 0.67 | | 0.92 |
| XYY | 0.94 | 0.85 | 0.84 | 0.78 | |

no GC correction / GC correction

**H3K9me2**

| | XY | XX | X0 | XXY | XYY |
|---|---|---|---|---|---|
| XY | | 0.95 | 0.77 | 0.75 | 0.84 |
| XX | 0.99 | | 0.82 | 0.73 | 0.78 |
| X0 | 0.72 | 0.72 | | 0.62 | 0.6 |
| XXY | 0.81 | 0.8 | 0.95 | | 0.76 |
| XYY | 0.95 | 0.94 | 0.86 | 0.92 | |

no GC correction / GC correction

**H3K9me3**

| | XY | XX | X0 | XXY | XYY |
|---|---|---|---|---|---|
| XY | | 0.78 | 0.68 | 0.64 | 0.8 |
| XX | 0.94 | | 0.75 | 0.41 | 0.62 |
| X0 | 0.79 | 0.81 | | 0.58 | 0.63 |
| XXY | 0.8 | 0.76 | 0.95 | | 0.71 |
| XYY | 0.94 | 0.92 | 0.9 | 0.9 | |

no GC correction / GC correction

**Table S3**. Pearson correlation coefficients for different ChIP experiments. Shown are correlation coefficients of H3K9me3 signal across different karyotypes. Correlation coefficients are calculated based on normalized ChIP-seq signal in 5-kb non-overlapping windows.

correlation of H3K4me3 signal across samples

| | XY | | | |
|---|---|---|---|---|
| XX | 0.898 | XX | | |
| XO | 0.926 | 0.963 | XO | |
| XXY | 0.866 | 0.816 | 0.832 | XXY |
| XYY | 0.893 | 0.879 | 0.89 | 0.748 |

H3K9me2 signal vs. H3K9me3 signal

Pearson correlation

| XY | XX | XO | XXY | XYY |
|---|---|---|---|---|
| 0.7356 | 0.9015 | 0.674 | 0.5335 | 0.7557 |

overlap of top 40% of 5kb windows

| XY | XX | XO | XXY | XYY |
|---|---|---|---|---|
| 0.6873 | 0.7166 | 0.5998 | 0.657 | 0.6866 |

Unspiked vs. spiked H3K9me3 replicates

Pearson correlation

| XY | XO | XXY | XYY |
|---|---|---|---|
| 0.7559 | 0.675 | 0.5665 | 0.8967 |

**Table S4**. GO categories of differentially expressed genes (***p<10-9, **p<10-6, *p<10-3).

| **GO term** | XX vs. XY | XX vs. XO | XX vs. XXY | XY vs. XO | XY vs. XYY |
|---|---|---|---|---|---|
| amide biosynthetic process | * | *** | ** | * | |
| amino sugar metabolic process | | *** | * | *** | |
| aminoglycan metabolic process | * | *** | | *** | |
| antibacterial humoral response | | | * | | |
| axoneme assembly | | | | | * |
| behavioral response to starvation | | | | * | |
| biosynthetic process | | | * | | |
| body morphogenesis | | | | * | |
| carbohydrate derivative metabolic process | | * | | * | |
| cellular amide metabolic process | * | *** | ** | * | |
| cellular biosynthetic process | | | * | | |
| cellular macromolecule biosynthetic process | * | * | ** | * | |
| cellular nitrogen compound biosynthetic process | * | | * | * | |
| cellular protein metabolic process | | | * | | |
| cellular response to heat | * | ** | ** | * | |
| cellular response to UV | * | | | | * |
| centrosome duplication | | ** | | | |
| chitin metabolic process | * | *** | * | *** | * |
| chitin-based cuticle development | | ** | * | *** | |
| coagulation | | * | | | |
| cold acclimation | | * | * | * | |
| cuticle chitin metabolic process | | | | * | |
| cuticle development | | ** | * | *** | |
| cytopasm organization | ** | ** | | | |
| defense response | * | | | | |
| defense response to bacterium | | * | | | |
| defense response to Gram-positive bacterium | ** | * | * | * | |
| dicarboxylic acid catabolic process | | * | | | |
| glucosamine-containing compound metabolic process | | *** | * | *** | |
| heat shock-mediated polytene chromosome puffing | | * | * | * | |
| hemolymph coagulation | | * | | | |
| hemostasis | | * | | | |
| humoral immune response | | | * | | |
| lipid catabolic process | * | | | | |
| macromolecule biosynthetic process | * | | * | * | |
| mating plug formation | | | | | * |
| melanin biosynthetic process | | * | | | |
| microtubule bundle formation | | | | | * |
| microtubule organizing center organization | | * | | | |
| mitotic spindle organization | | * | | | |

| | | | | | |
|---|---|---|---|---|---|
| multi-organism process | * | | | | * |
| multicellular organism reproduction | *** | | | *** | *** |
| negative regulation of female receptivity | | | | | * |
| nitrogen compound metabolic process | | | * | | |
| organic substance biosynthetic process | | | * | | |
| organonitrogen compound biosynthetic process | * | ** | ** | * | |
| organonitrogen compound metabolic process | * | *** | * | | |
| oxidation-reduction process | * | | * | | |
| peptide biosynthetic process | ** | *** | ** | * | |
| peptide metabolic process | * | *** | ** | * | |
| polytene chromosome puffing | | * | * | | |
| post-mating behavior | | | | | *** |
| regulation of rhdopsin mediated signaling pathway | | | | | * |
| reproduction | *** | | | ** | *** |
| reproductive behavior | | | | | * |
| response to bacterium | * | ** | * | * | |
| response to biotic stimulus | * | * | | | |
| response to chemical | | | * | | |
| response to cold | | * | | * | |
| response to external biotic stimulus | * | * | | | |
| response to heat | | * | ** | * | |
| response to organic substance | | | * | | |
| response to other organism | * | * | | | |
| response to pheromone | * | * | * | | * |
| response to temperature stimulus | | * | ** | * | |
| rRNA 2'-O-methylation | | * | | * | |
| rRNA metabolic process | | * | | | |
| rRNA methylation | | | | * | |
| rRNA processing | | * | | | |
| sensory perception of chemical stimulus | * | * | ** | ** | |
| sex differentiation | ** | ** | | | |
| sperm axoneme assembly | | | | | * |
| sperm chromatin condensation | | | | | * |
| sperm competition | | | | | * |
| sperm displacement | | | | | * |
| spermatogenesis | | | | | * |
| translation | ** | *** | ** | * | |
| vitellogenesis | *** | *** | | | |
